# Glia regulate local retinoic acid levels to specify neuronal specialisation for high-acuity vision

**DOI:** 10.64898/2026.02.27.708512

**Authors:** Manuela Lahne, Roxana Lungu, Madison Snorton, Takeshi Yoshimatsu, Ryan B. MacDonald

## Abstract

Neurons of the same type can be differentially tuned to perform distinct functional roles, yet how such specialisation arises during development remains poorly understood. Here, we show that in the zebrafish retina, heterogeneity within a single glial population generates local gradients of retinoic acid (RA) signalling that fine-tune neural function within a single neuronal type. The acute zone (AZ), a ventro-temporal region of the retina, is specialised for prey capture in the upper frontal visual field. In this region, UV cones exhibit enhanced light sensitivity through elongation of their outer segments (OS), the cellular compartment responsible for phototransduction. We found that *cyp26* genes, which encode RA-degrading enzymes, are expressed in Müller glia in a region-specific manner, and that this expression pattern closely matches the spatial variation in cone OS length. Inhibition of Cyp26 activity prevented OS elongation in the AZ. This effect is mediated by RA signalling, as direct activation of RA receptors using RARα agonists similarly reduced OS length in this region. Notably, inhibition of Cyp26 predominantly activated RA signalling in Müller glia rather than in cones, indicating that RA signalling regulates cone OS length in a cell-non-autonomous manner via Müller glia. Consistent with this model, blocking Müller glia genesis abolished cone specialisation across all regions of the developing retina. Together, these findings identify Müller glia as key regulators of cone functional specialisation through the spatial control of RA signalling.

## INTRODUCTION

Retinal functions are regionally tuned to process asymmetrical natural visual scenes and to serve visual field-specific behaviours [1,2]. Across many species, the frontal visual field is used for predation [3–6]. Correspondingly, the retinal region viewing the frontal visual field is often specialised for high-acuity vision, enabling fine spatial discrimination and detection of small objects [3,5,6]. In humans, this retinal region, the fovea, consists of a high density of cones in order to sample visual scene at higher spatial resolution [7]. Additionally, foveal cones are functionally specialised. They exhibit higher light sensitivity accompanied by elongated outer segment (OSs), the light sensing compartment of photoreceptors [8,9]. As cones are the most metabolically demanding cells in the retina and OSs account for a large fraction of this energy expenditure [10], the fovea and surrounding region, the macula, consume particularly high energy [10], which likely contributes to the macula’s vulnerability to degenerative diseases [11,12]. Understanding the mechanisms underlying these region-specific cone specialisations may provide important insights into the origin of macular diseases. However, it remains largely unclear how regional functional specialisations of the same type neurons is established during development.

Zebrafish is a diurnal teleost fish species with cone-rich retina and has been widely used to study photoreceptor development and model human eye diseases [13,14]. Similar to the human retina, zebrafish retinas display regional variation in cone OS length [3,15]. UV sensitive cone (PR4 in the vertebrate photoreceptor nomenclature [16]) OSs are longer in the horizontal midline areas than in the dorsal and ventral retina [3]. In particular, the acute zone (AZ), the retinal region for high-acuity vision, contains the longest cone OSs [3,15]. This region of the retina, which is located ventro-temporal region of the eye, features functional and structural specializations analogous to those of human foveal cones [3]. The AZ consists of a high density of UV cones, which mediates predation in zebrafish [3]. Accordingly, the light sensitivity of AZ UV cones is higher [3].

Retinoic acid (RA) is a morphogen that regulates multiple aspects of early stage eye development including eye morphogenesis, patterning, and photoreceptor differentiation [17–19]. In the developing retina, RA signalling activity is regionally restricted: high in dorsal and ventral regions and low in the horizontal midline areas or, in some species, in the central retina [20–22]. This spatial pattern arises from localized expression of RA - synthesizing and -degrading enzymes. RA synthesizing enzymes (RALDH1a2 and RALDH1a3) are expressed dorsally and ventrally [23–26], whereas RA-degrading enzymes (Cyp26a1 and Cyp26c1) are enriched along the horizontal midline area or central region [20,22,27]. In the developing chick retina, lack of RA signalling is critical for establishing the high acuity area, which is facilitated by regionalised Cyp26 expression [20]. This regional *cyp26a1* expression is also observed in the primate macula – the fovea and its surrounding region - where *cyp26a1* expression is specifically localized in Müller glia, suggesting a potential macula specific role in regulating RA signalling and high-acuity function [27].

Here, we explore whether RA signalling regulates morphological and functional specialisation of UV cones in the zebrafish AZ. We found that *cyp26* genes are specifically expressed in Müller glia along the horizontal midline area, with *cyp26a1* expression emerging during the developmental time window when UV cone OS elongation occurs. Pharmacological inhibition of Cyp26s activity during this period resulted in a loss of OS elongation in AZ UV cones. Activation of RA signalling via the RARα agonist, a RA receptor, similarly abolished OS elongation and reduced light sensitivity in UV cones, indicating that the effect of Cyp26 inhibition is mediated by increase in RA signalling. These findings suggest that local degradation of RA by *cyp26* in Müller glia is essential for establishing regional specializations of cones. We further tested whether inhibition of Cyp26 in Müller glia increases RA signalling in nearby UV cones. To our surprise, we found that Cyp26 inhibitor activates RA signalling primarily in Müller glia across the retina rather than in cones. This cell type-specific activation of RA signalling was also observed after RARα agonist treatment, suggesting that RA regulates cone specializations in a cell non-autonomous manner through Müller glia. Consistent with this, blocking Müller glia genesis in the retina eliminated cone OS specialization. Together, these results identify Müller glia as key regulators of regional neuronal specialization through local modulation of RA signalling.

## Material and Methods

### Zebrafish maintenance and handling

All procedures for experimental protocols were approved by the UCL AWERB (Animal Welfare and Ethical Review Body) in addition to the U.K. Home Office (PPL no. PP2133797 held by RBM) and Washington University in St Louis Institutional Animal Care and Use Committee guidelines. Zebrafish were raised and maintained under a 14:10 hour light:dark cycle at 28.5 °C.

Zebrafish embryos obtained from pair or group matings that were designated for experiments were treated with 100 µM PTU (1-phenyl 2-thiourea, Sigma-Aldrich) dissolved in E3 medium (5 mM NaCl, 0.17 mM KCl, 0.33 mM CaCl2, 0.33 mM MgSO4) starting at around 8 hours post fertilisation (hpf). The E3 medium containing PTU was exchanged daily. Embryos/larvae were staged according to established anatomical criteria (Kimmel et al., 1995). The following zebrafish lines were used in this study: *Tg(tp1bglob:VenusPest)^s940^* [28] referred to as *Tg(tp1:Venus)*, *Tg(TP1glob:eGFP-CAAX)^u911^*[29] referred to as *Tg(TP1:GFP-CAAX))* and *Tg(CSL:mCherry)^jh11^* [30] to identify MG, *Tg(-5.5opn1sw1:EGFP)^kj9^*referred to as *Tg(opn1sw1:GFP)* to mark UV cones [31], and *Tg(12xrare-ef1a:EGFP)^sk71^ (referred to as Tg(RARE:GFP))* as a reporter of RA signalling [32]. *Tg(opn1sw1:sypb-GCaMPf)^uss101^*to express synaptically localized calcium sensor, SyGCaMP6f [3].

### Drug treatments

Zebrafish larvae that were treated with PTU from 8 hpf were immersed in PTU-containing E3 medium that was either supplemented with the 2.5 µM R115866 (Cyp26 inhibitor, Merck Life Science Limited) [33], 50 nM AM580 (a retinoic receptor α agonist, Cayman Chemical) [32], 250 nM BMS961 (RARγ agonist, Tocris) [34] or the vehicle control DMSO (Life Technologies) at 72 hpf. The E3 medium containing PTU and the corresponding drugs were exchanged at 96 hpf.

To disrupt Müller glia genesis, dechorinated zebrafish larvae were exposed to 21 µM DAPT (N-[N-(3,5-Difluorophenacetyl)-L-alanyl]-S-phenylglycine t-butyl ester, Merck Life Science Limited) or DMSO in PTU-containing E3 medium from 45 to 120 hpf [35]. At 72 hpf, DAPT- or DMSO-treated zebrafish were additionally exposed to either 50 nM AM580 or DMSO (see schematic in Fig. 10A) and solutions were replenished at 96 hpf. In a subset of experiments, zebrafish larvae were treated with 21 µM DAPT or DMSO in E3 medium supplemented with PTU from 72 to 120 hpf and solutions were exchanged at 96 hpf (Fig. 10F). Larvae were terminally anesthetised in MS-222 (Ethyl 3-aminobenzoate methanesulfonate, Sigma-Aldrich) at 120 hpf and then fixed in 4 % paraformaldehyde (PFA, Fisher Scientific Ltd) in PBS for 30 min or 1 hour at room temperature (RT) for subsequent use in immunocytochemical experiments or to perform HCR (hybridisation chain reaction) labelling, respectively. For RNA isolation, zebrafish larvae were fixed in 4 % PFA for 30 min at 4 °C.

### Immunohistochemistry

Immunolabelling was performed on isolated eyes according to [3] with minor adjustments. In detail, following fixation in 4% PFA, larvae were washed three times in PBS for 10 min. Subsequently, the eyes were removed and incubated in PBS containing 0.1% TritonX (PBST, Sigma-Aldrich) overnight at 4 °C. Eyes were blocked in PBS supplemented with 10% goat serum, 1% bovine serum albumin and 0.1% TritonX for 2 hours at RT and then incubated in primary antibodies for 2 days at 4 °C. Primary antibodies to GFP (1:1000, chicken, GeneTex) and UV opsin (1:1000, rabbit, Kerafast) were diluted in blocking solution that additionally contained 2 % DMSO and 1% Tween20 (Sigma-Aldrich). The eyes were washed four times in PBST for one hour each at RT. Subsequently eyes were incubated with goat anti chicken Alexa fluor-488 (1:1000, Thermo Fisher Scientific), goat anti rabbit Alexa fluor-647 (1:1000, Thermo Fisher Scientific) secondary antibodies in blocking solution with DMSO (2%) and Tween20 (1%) added and incubated for two days at 4 °C. The eyes were washed four times in PBST before they were mounted in 1% low melting point agarose (Sigma-Aldrich) dissolved in PBS on a glass bottom dish (Nunc, Scientific Laboratory Supplies) for confocal imaging.

### Hybridisation chain reaction *in situ* hybridisation

Hybridisation chain reaction (HCR) *in situ* hybridisations were carried out according to [36] with minor modifications: PFA-fixed *Tg(tp1bglob:VenusPest)s940* or *Tg(RARE:GFP)* larvae (48, 60, 72 and 120 hpf) were incubated in 20 µg/ml Proteinase K (Roche) diluted in PBS for 20 min (48, 60 hpf) or 25 min (72, 120 hpf) at RT, then washed twice briefly in PBS prepared in DEPC (diethyl pyrocarbonate) water followed by fixation in 4 % PFA for 20 min at RT. Subsequently, larvae were washed three times 5 min in PBS containing 0.1 % Tween20 and then incubated for 30 min in pre-warmed hybridisation buffer (Molecular Instruments). Larvae were exposed to HCR probes (Table 1) diluted at 16 nM in hybridisation buffer for two days at 37 °C. Larvae were washed in probe wash buffer (Molecular Instruments) four times for 15 min each at 37 °C and then in 5x SSCT (Sodium chloride, Sodium citrate containing 0.1 % Tween20) for 5 min at RT. 12 pmol of hairpin 1 and 2 of the HCR B1-647-conjugated or B2-546-conjugated amplifier sets (Molecular Instruments) were incubated individually at 95 °C and snap cooled before they were mixed in amplification buffer (Molecular Instruments). Larvae were exposed to amplifiers for two days at RT and subsequently washed four times for 5 min each in SSCT buffer before they were mounted on glass bottom dishes in 1 % low melting point agarose for confocal imaging.

**Table 1:**
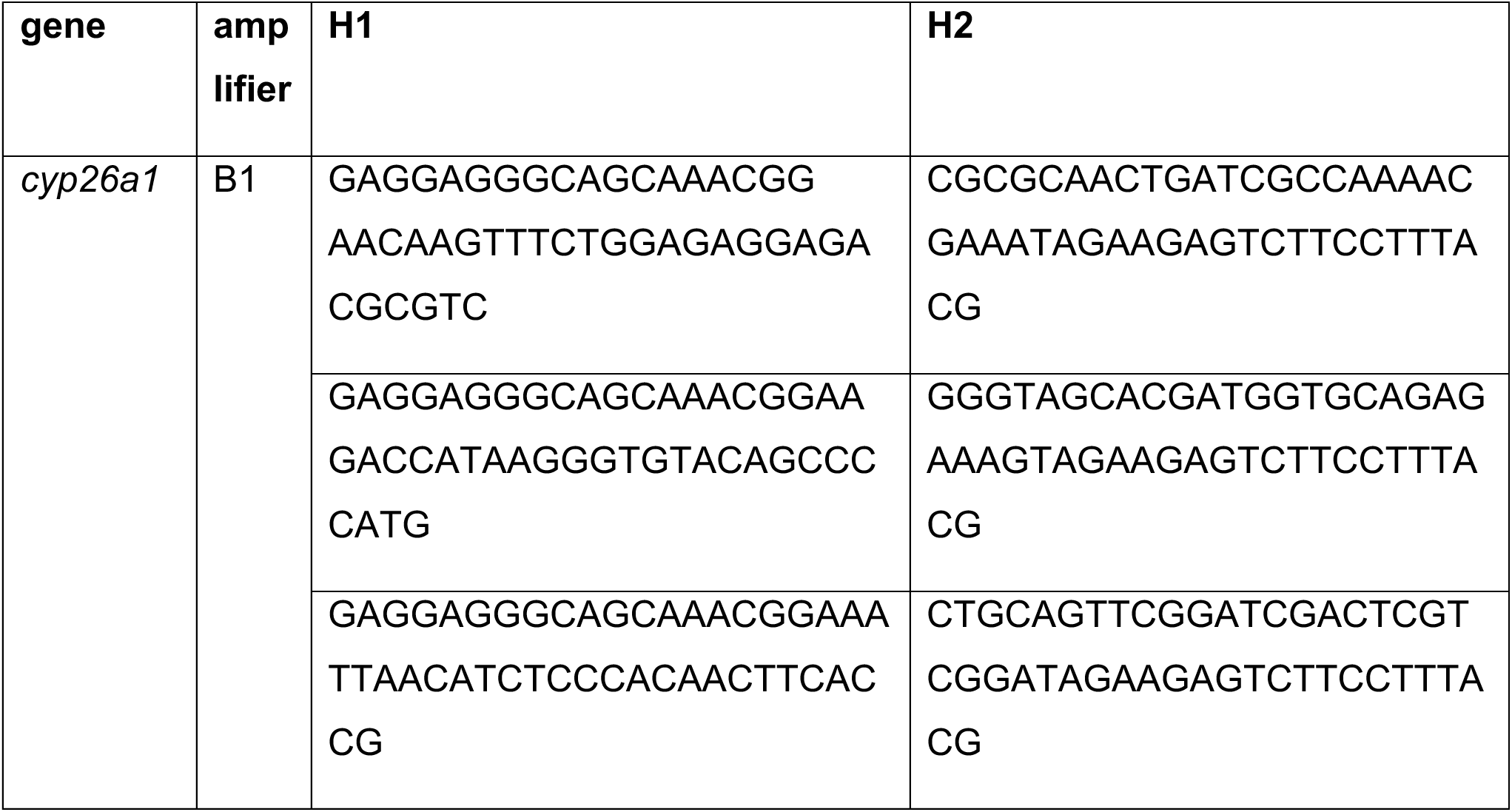
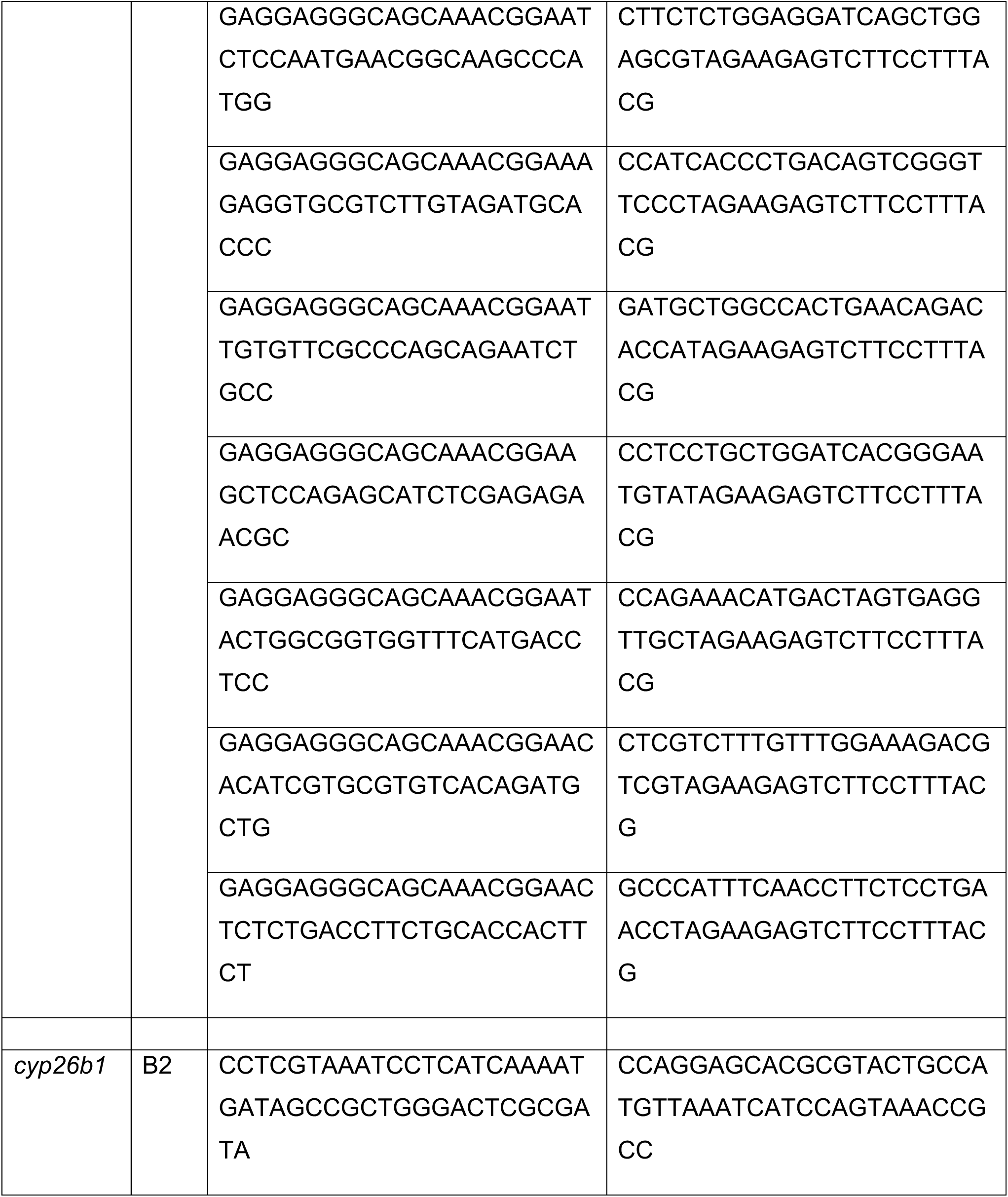
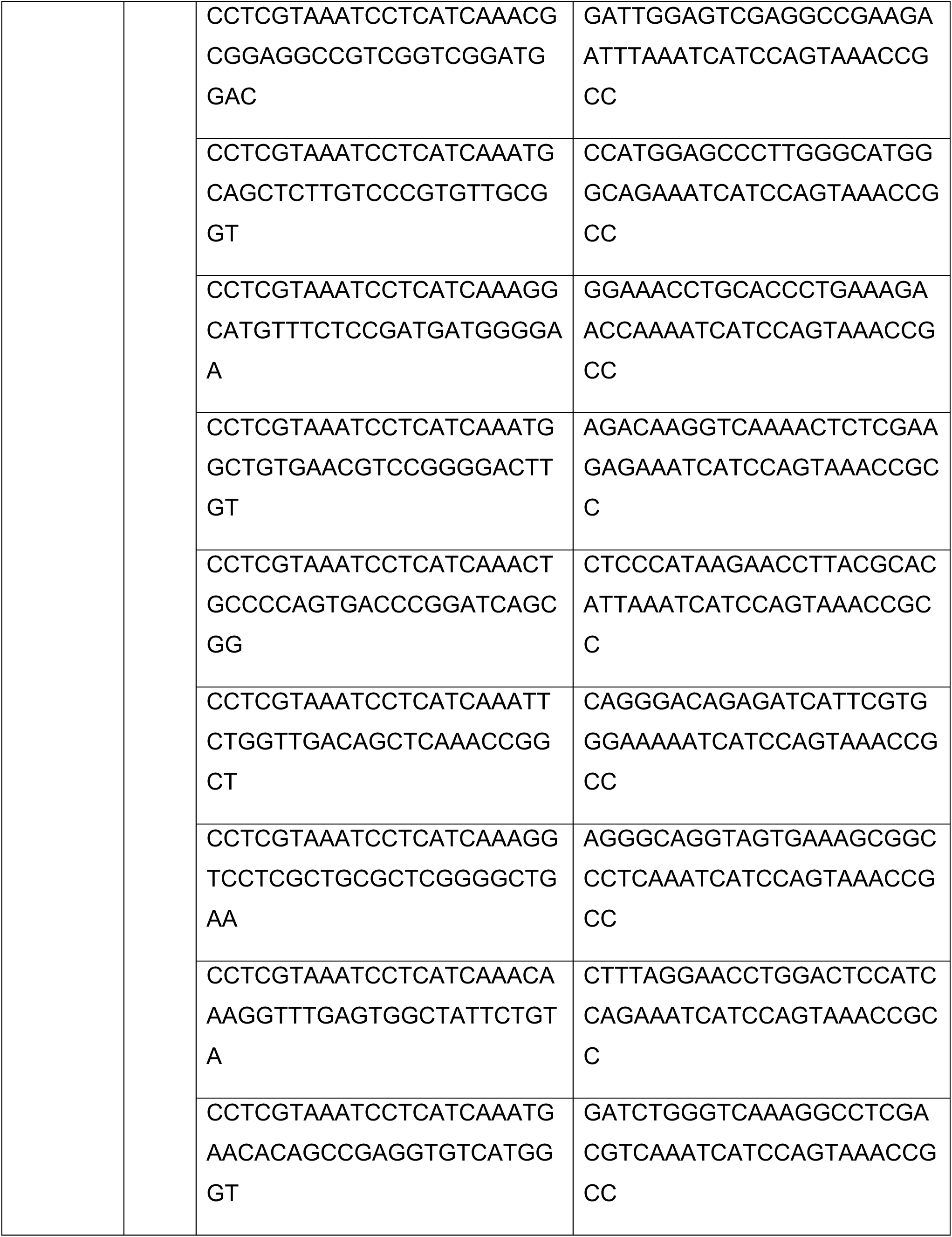
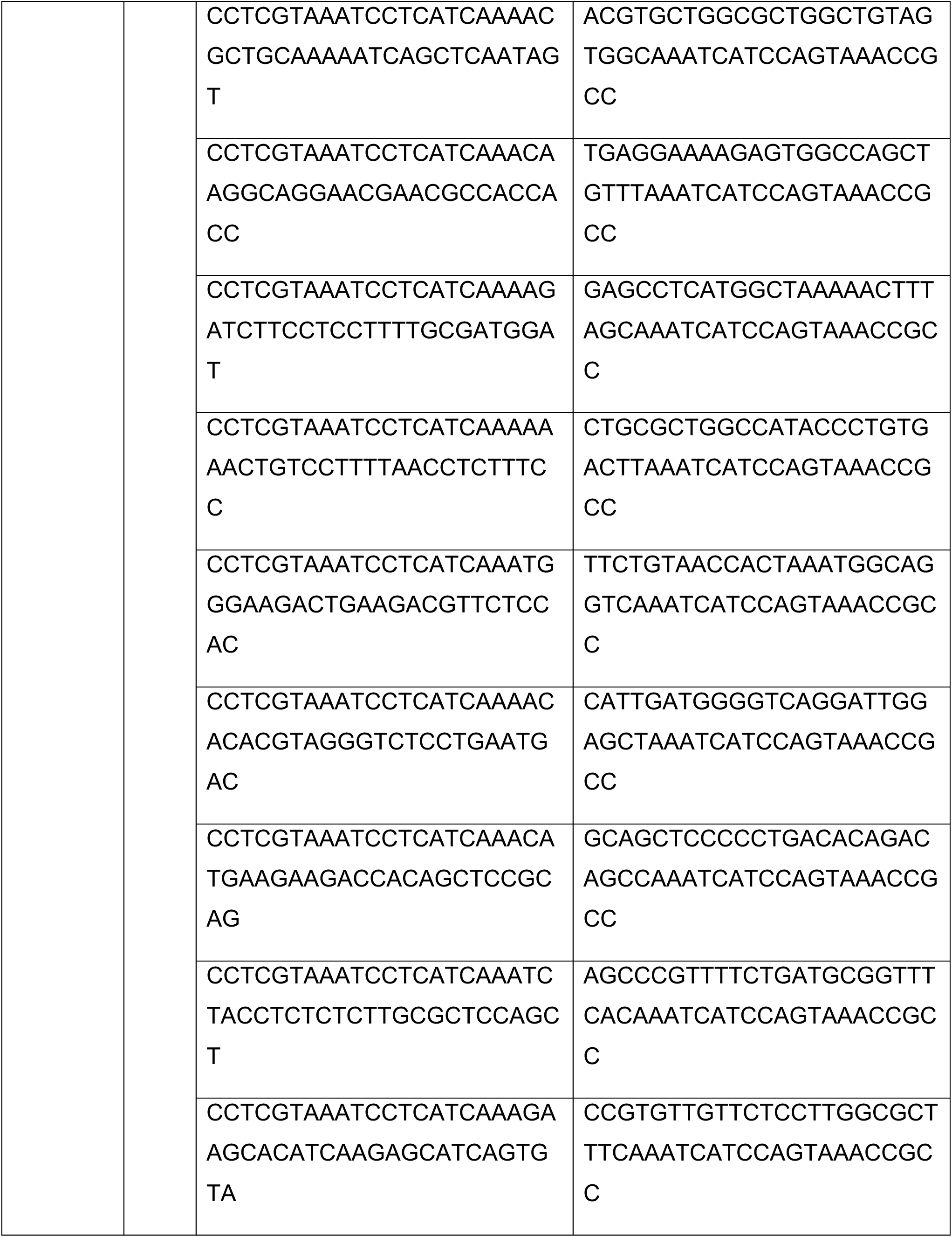
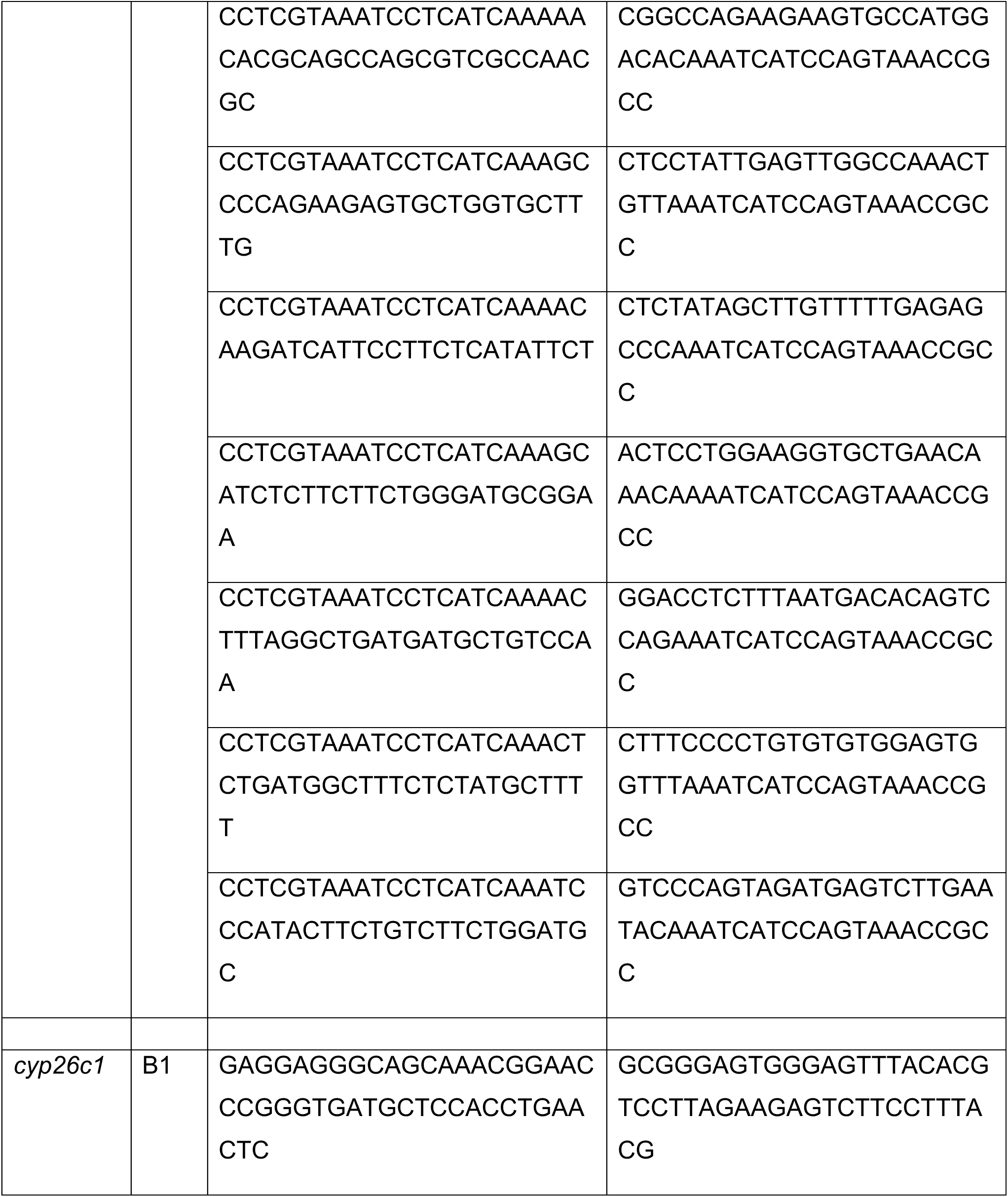
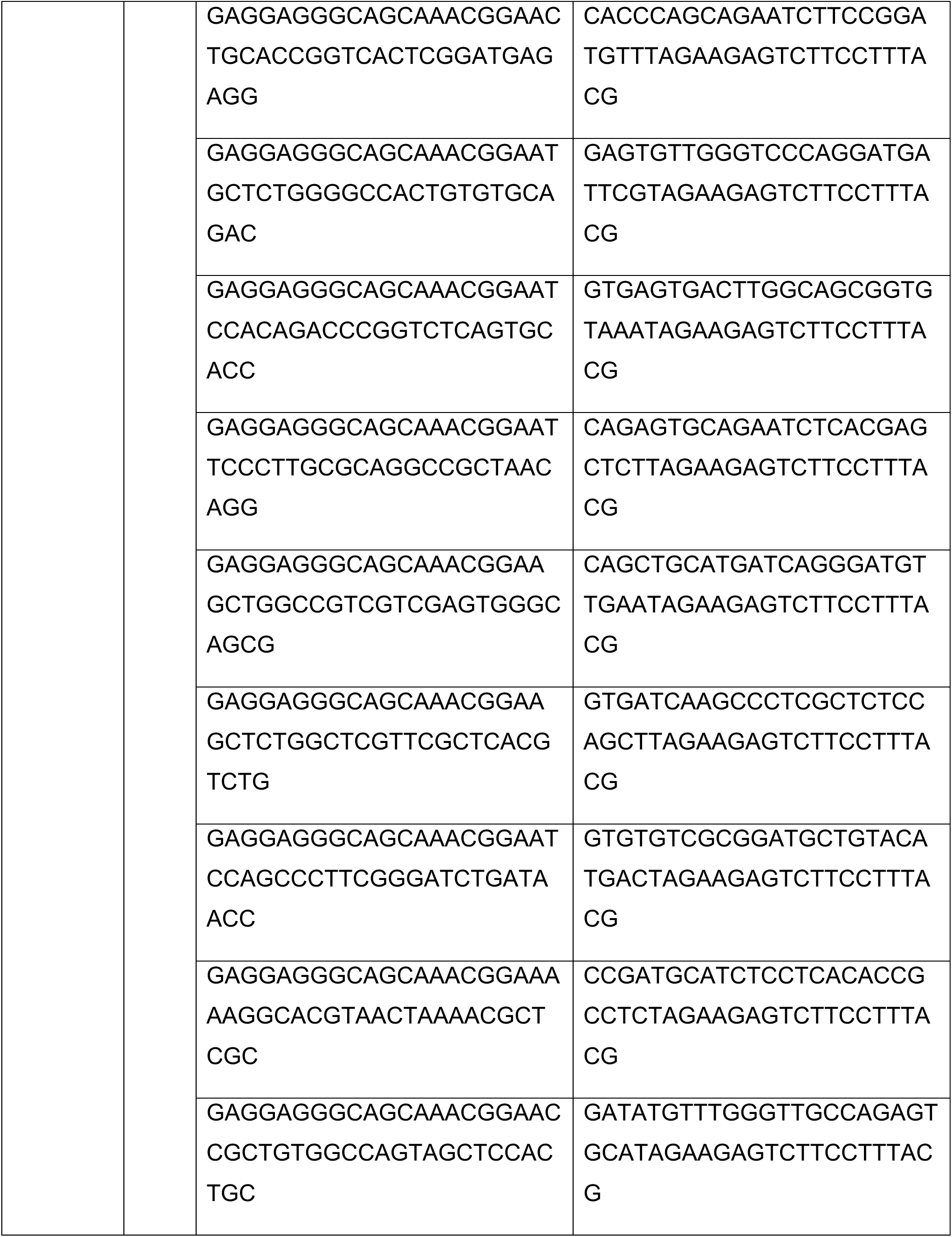
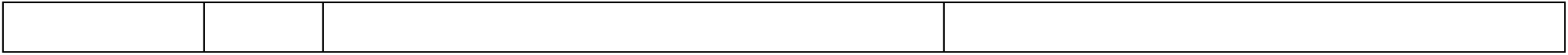
List of HCR probes for *cyp26* and *rar* genes.

### Confocal microscopy

Z-stacked images (1024x1024) of the AZ, dorsal and nasal regions of 72 to 120 hpf immuno-labelled larval eyes were acquired using a Zeiss LSM900 confocal microscope equipped with an Airyscan 2 unit, 40x long distance C-Apochromat water immersion objective (NA 1.1) and 405, 488, 561 and 640 nm laser lines [29]. To image at a consistent depth, the z-stacks were centred 30 µm from the top of the eye surface, which was based on the observation of *Tg(opn1sw1:GFP)-*positive cells or other cell-type specific markers. For a subset of images, subregions of the ONL and subretinal space were acquired by 4Y multiplex airyscan imaging 15 µm z-stacks (2984x1200) at optimal z-step size using the 40x long distance C-Apochromat water immersion objective (NA 1.1) at 1.3 zoom. Representative confocal single z-plane or a maximum projection of z-stack images were chosen for presentation purposes.

RA agonist or DAPT-treated live *Tg(RARE:GFP)* zebrafish larvae or double crosses with *Tg(CSL:mCherry)^jh11^* were anaesthetised in 65 µM MS-222 at 96 hpf and mounted in 1 % low melting point agarose in E3 medium in a lateral orientation with the eye facing the coverslip of the glass bottom dish and maintained in E3 medium containing MS-222. A 20x Plan-Apochromat objective (NA 0.8, M27) was used to acquire 80 µm z-stack images (512 x512) with a z-step of 1 µm to capture the majority of the thickness of the eye. For presentation purposes, single z-plane sections from the centre of the retina are shown.

HCR-labelled *Tg(tp1:Venus) or Tg(RARE:GFP)* zebrafish larvae were whole-mounted in a lateral orientation with the eye/lens facing the coverslip of the glass bottom dish. Z-stack images of 80 (48, 60 hpf) or 90 µm (72, 120 hpf) at a step-size of 1 µm were acquired with a 40x long distance C-Apochromat water immersion objective (NA 1.1) at 0.5 zoom using 488, 561 and 640 nm laser lines. Typically, one-tile images of 1024x1024 were acquired, however, for 120 hpf embryos 2 or 4-tile z-stack images of 1024x1024 each were acquired to allow visualising the labelling of the entire eye. For presentation purposes, maximum intensity projections of images spanning 32 µm (48, 60 hpf) or 42 µm thickness (72, 120 hpf) were generated. The chosen number of z-planes used to generate the maximum projection was increased due to the increased eye size at later stages.

### Image analysis

The length of UV cone OSs was measured in the AZ, dorsal and nasal regions of 16 cells each for drug- and DMSO-treated immunolabelled eyes. The segmented line tool in Image J was placed on the UV cone OS and the length was determined using the measurement function. The UV cone OS length of the 16 different cells was averaged to yield one value per region per eye (n>11 larvae analysed per treatment). For statistical analysis of multiple comparisons, ANOVA and a Tukey’s post-hoc test were performed.

The number of *RARE:GFP*-positive cells was either determined based on their location within the retina or on immuno-cytochemical co-labelling with cell type specific markers. Cells within the AZ, nasal, dorsal and ventral were counted in a region that spanned a 100 µm width and 120 µm height. Note, ventral RPE cells were not quantified as these were often overexposed to visualise the other less brightly labelled cell types. For statistical analysis of multiple comparisons, ANOVA and a Tukey’s pos-thoc test were performed.

### RNA isolation and quantitative reverse transcriptase polymerase chain reaction (qRT-PCR)

In a subset of experiments, drug-exposed larvae (AM580, BMS961) or their DMSO controls at 120 hpf were terminally anesthetised, then fixed in 4 % PFA for 30 min at 4 °C and washed twice in PBS before whole eyes were removed from the larvae in PBS. Eyes were transferred into Eppendorf tubes and the PBS was removed prior to quick-freezing the tissue at -80 °C. RNA was isolated using the FFPE RNAeasy kit according to manufacturer’s recommendations (Qiagen). RNA was transcribed into cDNA using qScript™ cDNA SuperMix (Quantabio). Expression levels of RA target genes were assessed using the QuantStudio 6 qPCR machine on samples that were amplified using the SYBR Select Master Mix (Thermo Fisher) in combination with the primers listed in

Table 2. The delta delta Ct method with actin as a house keeping gene was employed to determine expression differences between drug-treated and control samples.

**Table 2:**
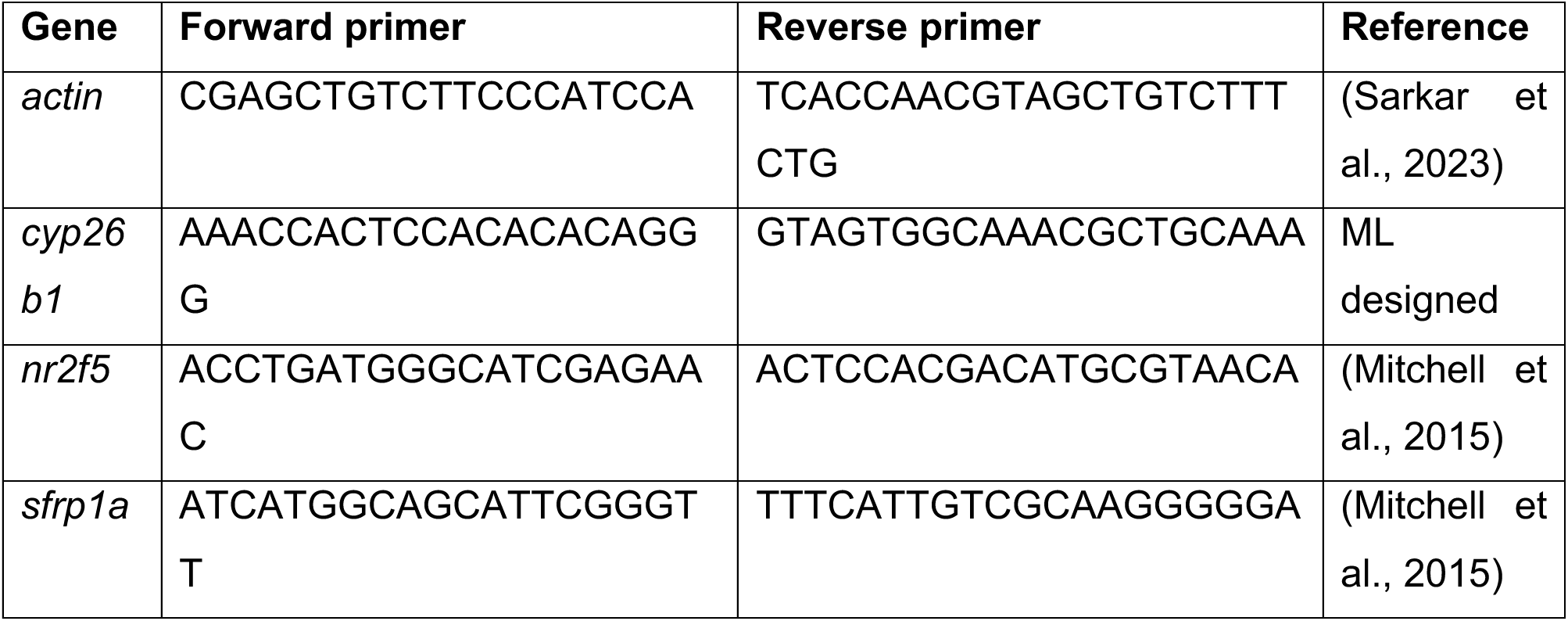
Primers used for qRT PCR.

### Two-photon UV cone calcium imaging and light stimulation

Two-photon imaging of UV cone calcium responses was performed on a custom-build microscope equipped with a mode-locked Ti:Sapphire laser (Spectra-physics MaiTai DeepSee) tuned to 950 nm, resonant-galvo scanner (MBF Bioscience), and a water immersion objective (Zeiss W Plan-Apochromat 20x/1.0 DIC UV-VIS). We used the transgenic line *Tg(opn1sw1:SyGCaMP6f)* that expresses a synaptically targeted calcium sensor in UV cones. Fish were immobilized by injection of α-bungarotoxin (1 nL of 2 mg/ml; Tocris, Cat.2133) into the space between the skull and the eye. They were mounted on the side in 1.5% low melting point agarose (Sigma-Aldrich, Type VII-A, Cat. A0701) in a glass bottom chamber and submerged in fish water. Fluorescent from SyGCaMP6f was isolated by a bandpass filter (Thorlabs FBH520-40) placed in front of a photomultiplier detector (Hamamatsu H10770A-40). For image acquisition, we used ScanImage (MBF Bioscience). Typical recording configurations were 256 x 60 pixels (1 ms per line, 16.7 frames per second). Fish were mounted on the side in a glass bottom chamber, imaged from the top, and light stimulation was delivered from the bottom by light emitting diodes (LEDs) (Roithner, UVLED365-11E, 365 nm). LED light was filtered using a dichroic mirror (Chroma, T425lpxr) and synchronized with the scan overshoot at 1 kHz. The intensity of the LED light was 8 nW/mm^2^ (∼1.5x10^4^ photons/sec/µm^2^) at the fish eyes.

## Results

### Regional difference in OS growth underlies variations in the UV cone OS length

While UV cone OS development was previously described [37], it is currently unknown how regional OS length differences are established during retinal development. We assessed the timing of UV cone OS growth in the AZ, dorsal and nasal regions of the retina (Fig. 1A) from cone genesis at 72 hpf until 120 hpf when larval zebrafish perform predation (Fig. 1B). To measure the UV cone OS length (Fig. 1C), we used *Tg(opn1sw1:GFP)* zebrafish immunolabeled with UV opsin (Fig. 1D). At 72 hpf, UV cones are fate-specified as evidenced by the presence of UV opsin in their OSs. At this timepoint, their OS length was of similar length in all regions assessed (Fig. 1D,E). However, UV cone OS length began to differentially increase in each region by 96 hpf and by 120 hpf (Fig. 1D,E). The regional UV cone OS length differences were evident with those UV cones in the AZ and nasal regions possessing significantly longer OSs compared to the dorsal retina (Fig. 1D,E). Thus, differential growth underlies regional differences in UV cone OS length and that specialisation of UV cone OS length in the AZ is already apparent by 120 hpf in the zebrafish retina.

**Figure 1.**
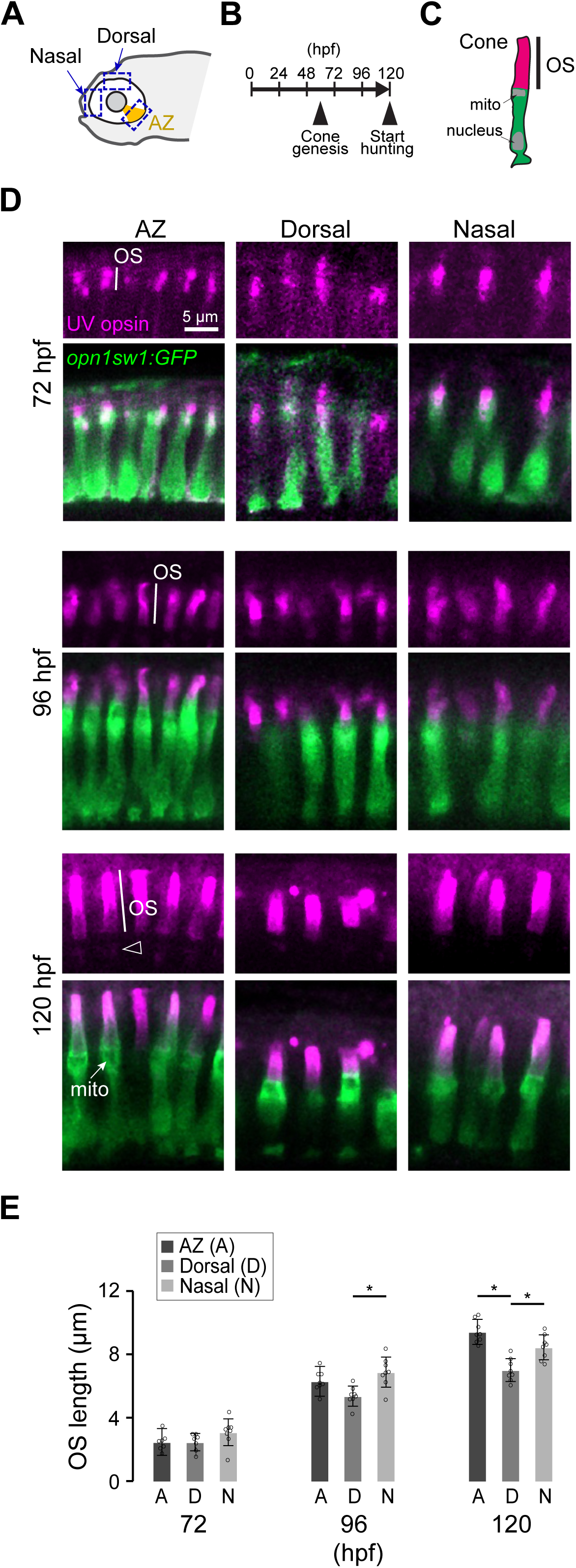
Regional differences in photoreceptor outer segment (OS) length are evident by 120 hours post-fertilization (hpf). **A**, Schematic of the zebrafish head positioned laterally. The orange area in the eye indicates the acute zone (AZ). Dashed blue rectangles indicate the imaged regions. **B**, Timeline of cone development and visual behavior in zebrafish. **C**, Schematic of a photoreceptor and its cellular components: the OS, mitochondria (mito), and nucleus. A black line indicates the length of the OS, which was measured in (D). **D**, Confocal images of UV cones (Green) and UV opsin immunostaining (magenta) in the *Tg(opn1sw1:GFP)* transgenic line. Note weak UV opsin staining is seen outside of the OS (open arrowhead), beneath mitochondoria (arrow) at 120 hpf. **E**, A bar graph of mean UV cone OS length at 72, 96 and 120 hpf for the AZ, dorsal and nasal regions. n= 7 (AZ), 7 (dorsal), 8 (nasal) at 3 dpf, and n=8 for all other ages and regions. Open circles: data from individual fish. Error bars: Standard deviation. One-way ANOVA plus Tukey’s post-hoc test, *: p < 0.05.

### Spatial RA signalling activity pattern inversely correlates with UV cone OS length

Seeking mechanisms that regulate cone outer segment length, we identified that RA signalling activity varies across the retina in a pattern corresponding to the outer segment length differences [21]. RA acts on a heterodimeric retinoic acid receptor (RAR) and retinoid X receptor (RXR) complex, which binds to retinoic acid response elements (RARE) within promoters or other gene regions (e.g., introns) to facilitate target gene expression or suppression upon RA binding [17]. We quantified RA signalling activity across the retina using *RARE:GFP* transgenic line [38]. This transgenic contains 12 repeats of RARE sequences followed by minimal promoter to express GFP gene in a RA signalling dependent manner (Fig. 2A,B). Comparison of RA signalling activity and UV cone OS length (data from [3]) at 8 dpf revealed a tight inverse correlation between these two (Fig. 2C,D), indicating that RA is a prominent candidate for a signalling pathway regulating cone OS length.

**Figure 2.**
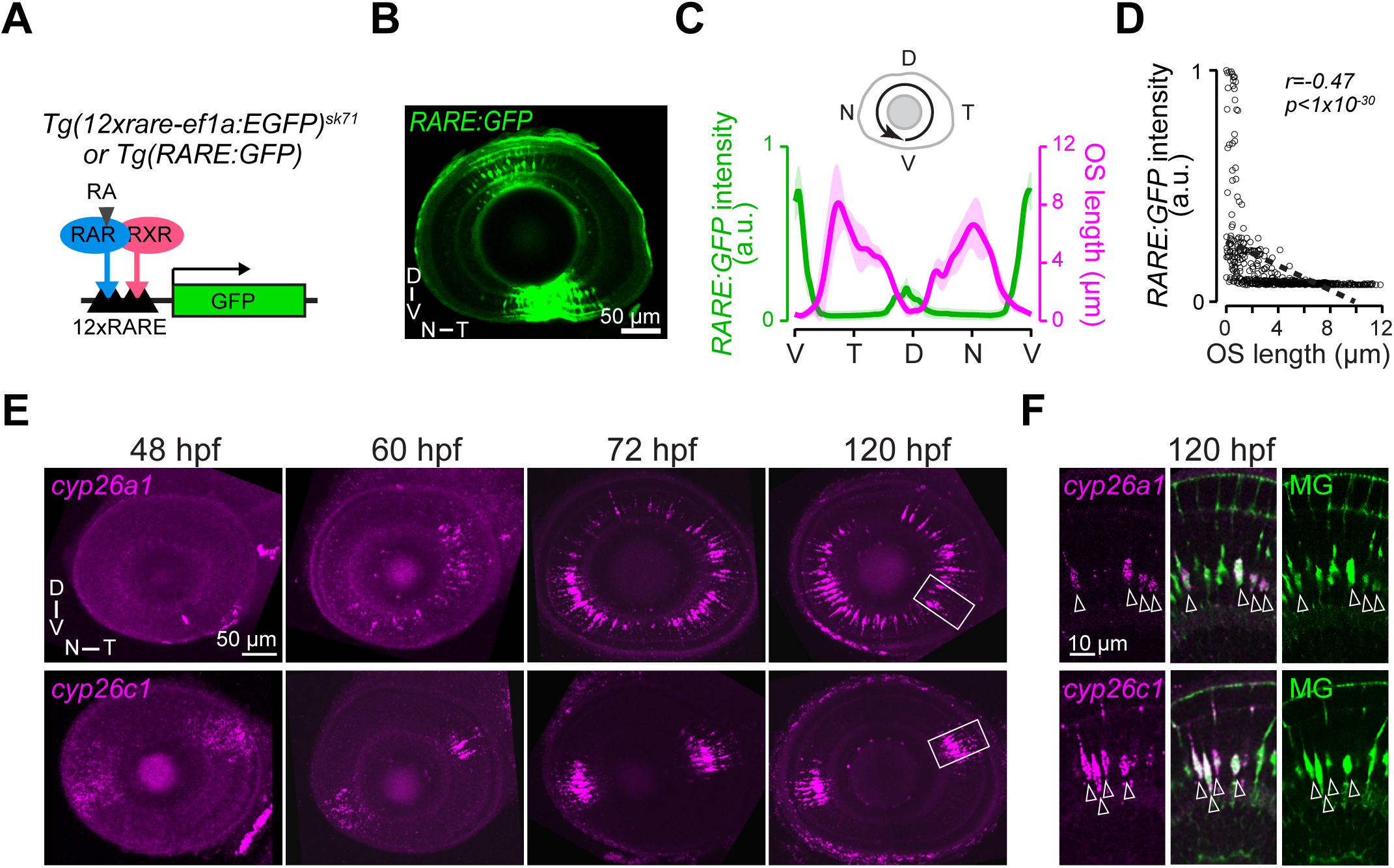
*cyp26a1* and *c1* are expressed by Müller glia in the retinal areas with low Retinoic acid signalling activity which inversely correlates with regional differences in UV cone OS length. **A**, Schematic of retinoic acid signalling reporter construct in *Tg(12xrare-ef1a:EGFP)^sk71^*transgenic zebrafish. Retinoic acid receptor (RAR) and Retinoid X receptor (RXR) complex bind to Retinoic Acid Response Element (RARE) and transactivate downstream gene expression upon binding to Retinoic acid (RA). The reporter construct contains twelve repeats of RARE sequence followed by a minimum promoter ef1a and GFP coding sequence. **B**, A confocal image of a *Tg(RARE:GFP)* transgenic eye at 120 hpf. **C**, Line profiles of GFP intensity (green, arbitrary unit, a.u.) and UV cone OS length (magenta) across the eye, measured from the ventral region counter-clock direction as shown in the schematic. The data for UV cones OS length is from [3]. n=3 for *RARE:GFP* intensity. Line: mean, shades: standard deviations. **D**, *RARE:GFP* intensity as a function of UV cone OS length. Open circles: data from individual cone. Dashed line: linear trend line, *r* and *p*: Peason correlation coefficient and *p*-value. **E**, Confocal images of HCR in situ hybridisation for *cyp26a1* (top) and *cyp26c1* (bottom) at different time points during eye development. **F**, The *cyp26a1* and *cyp26c1* mRNAs (magenta) co-localise with Müller glia (MG) (open arrowheads), which are labeled (green) by Venus expression in *Tg(tp1:Venus)*. The regions displayed are indicated as white rectangles in **E**. D, dorsal; N, nasal; T, temporal; V, ventral.

### *cyp26a1* and *cyp26c1* are specifically expressed in Müller glia in the developing retina

We next investigated how the regionally restricted RA signalling activity arise. Previous studies showed RA synthesizing enzymes (*raldh1a2* and *raldh1a3*) are expressed in dorsal and ventral regions, respectively, in the developing zebrafish eye [25,26]. However, the expression patterns of *cyp26* genes in the developing eye in zebrafish and their contribution to shape the spatial RA signalling activity are unknown. Thus, we examined the regional expression patterns of *cyp26a1*, *cyp26b1* and *cyp26c1* in the zebrafish retina at different developmental timepoints (48, 60, 72 and 120 hpf) using HCR *in situ* hybridisation. The expression of *cyp26a1* was first observed from 60 hpf in the temporal and to a lesser extend in the ventral region of the retina and became predominant in the nasal and temporal retina from 72 to 120 hpf (Fig. 2E). Co-labeling with Müller glia using *Tg(tp1:VenusPest)* in which fluorescent protein VenusPest is expressed in Müller glia during early development [35] showed that *cyp26a1* is expressed in a subset of Müller glia (Fig. 2F). In contrast, cyp*26c1* expression is seen already at 48 hpf and spatially restricted to a small area in the ventro-nasal and dorso-temporal regions of the retina (Fig. 2E). This expression pattern persists at subsequent timepoints and resides in *cyp26a1* positive areas (Fig. 2E). Co-labeling with Müller glia in *Tg(tp1:VenusPest)* revealed that *cyp26c1* is also specifically expressed in Müller glia (Fig. 2F). No significant expression levels were detected for *cyp26b1* (Supplementary Fig. 1) as previously reported [39]. These findings indicate that *cyp26a1* along with *cyp26c1* expression closely resembles the regions with low RA signalling activity (Fig. 2E,F) and suggests that *cyp26a1* and *c1* in Müller glia may play critical role in establishing regionally restricted RA signalling activity.

### Cyp26 inhibition leads to regional UV cone OS shortening in the zebrafish retina

Having observed that *cyp26a1* and *cyp26c1* expression correlates with low RA signalling regions, which are also associated with longer UV cone OS regions (Fig. 2C), we examined the effects of disrupting Cyp26a1 and c1 function during OS growth. To disrupt Cyp26 function, embryos were exposed to the Cyp26 inhibitor R115866 (also known as talarozole) [33] from 72 to 120 hpf (Fig. 3A,B). The UV cone OS length was significantly shorter in the AZ of R115866-treated retinas compared to vehicle controls (Fig. 3C,D). In contrast, dorsal and nasal UV cone OS lengths were not statistically different between R115866-exposed and control retinas (Fig. 3C,D). We also measured the ratio of the UV cone OS length of the AZ or the nasal retina and the dorsal retina. These measurements compare relative OS growth of the AZ and the nasal OSs by normalising OS length against dorsal OS within individual retinas. In controls the ratio of the AZ and dorsal (AZ/Dorsal) and that of nasal and dorsal (Nasal/Dorsal) UV cone OS lengths were above 1 (Fig. 3E) as the AZ and nasal UV cone OSs are longer than those in the dorsal retina (Fig. 3E). While the nasal to dorsal ratio was unaffected in R115866-exposed retinas (Fig. 3E), the dorsal to AZ ratio was significantly decreased to below 1 in R115866-treated retinas, indicating that the UV cone OSs in the AZ are shorter than those in the dorsal region. These findings show that disrupting Cyp26 function, which is likely accompanied by accumulation of RA in the developing retina, abolishes elongation of OS in the AZ, suggest that low RA levels are critical for cone OS specialisation in the AZ.

**Figure 3.**
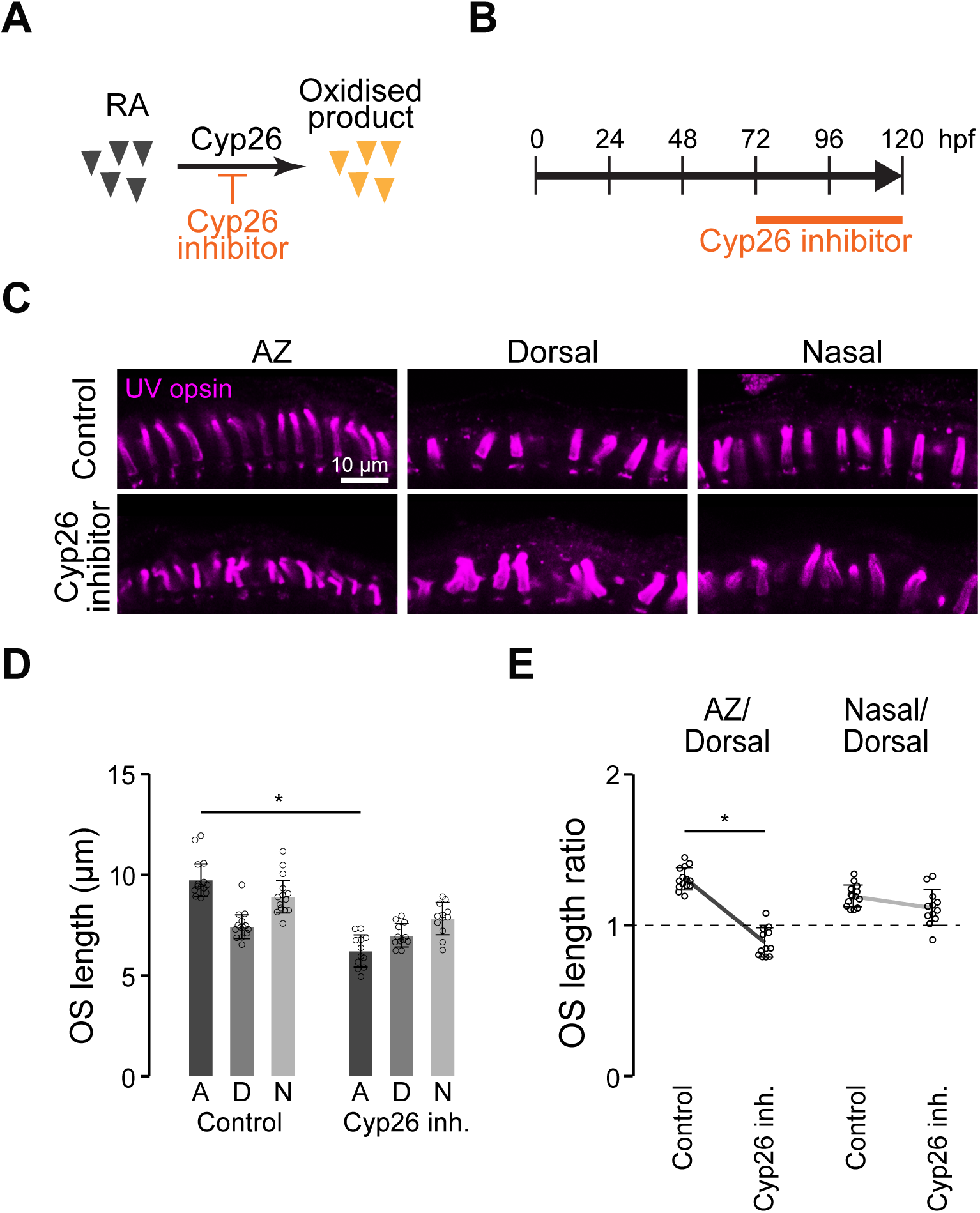
Cyp26 inhibition leads to shorter OS in the AZ. **A**, Schematic illustrating Cyp26-mediated degradation of retinoic acid is blocked by Cyp26 inhibitor leading to increased retinoic acid levels. **B,** Experimental timeline of drug application from 72 to 120 hpf. **C**, Confocal images of UV opsin in the photoreceptor layer in the AZ, dorsal and nasal retinas labelled by immunostaining at 120 hpf. Zebrafish were treated with control vehicle or a Cyp26 inhibitor (inh.), R115866, from 72 to 120 hpf. **D**, UV cone OS length in the AZ, dorsal and nasal regions of the retina at 120 hpf. **E**, Comparisons of the ratios of the UV cone OS length of the AZ or nasal regions against the dorsal region with and without R115866. Bars and Lines: mean. Error bars: standard deviation. Open circles: data from individual fish. n=14 (Control) and 12 (Cyp26 inhibitor, inh.) One-way ANOVA plus Tukey’s post-hoc test, *: *p* < 0.05.

### Cyp26 inhibition upregulates RA signalling in the retina in non-UV cone cells

To determine whether inhibition of Cyp26 leads to accumulation of RA we used Tg(*RARE:GFP)* transgenic zebrafish larvae. *RARE:GFP* zebrafish larvae were either exposed to a Cyp26 inhibitor, R115866, or vehicle from 72 to 120 hpf (Fig. 4A) and the eyes were imaged at 120 hpf (Fig. 4B,C). In vehicle controls, GFP was brightly expressed in the ventral retina and at a lower level in a subset of cells in the dorsal retina (Fig. 4B,C) similar to previously described data for earlier developmental timepoints at 56 hpf [21]. In contrast, exposure of larvae to R115866 resulted in increased GFP expression throughout the entire retina (Fig. 4B,C). Having observed an effect of R115866 on UV cone OS growth in the AZ we further examined GFP expression in the outer nuclear layer (ONL) in the different retinal regions. In controls, a few ONL cells expressed GFP in the ventral region, while only an occasional GFP*-*positive cell was present in the dorsal and none in the AZ or nasal ONL (Fig. 4D). Among GFP-positive cells in the ONL, only 0.83 ± 0.37 cells/retina in the ventral ONL co-labelled with UV opsin (Fig. 4E). Most GFP-positive cells in the ventral ONL are red and green cones that are marked by zpr-1 antibody (Fig. 4F). Surprisingly, R115866 did not significantly induce expression of GFP in ONL cells in any of the regions (Fig. 4D). The same was observed when GFP expression was assessed in specific cone subtypes (Fig. 4E,F). This data indicates that RA does not activate the down-stream signalling in UV cones in the AZ, suggesting that RA regulates UV cone OS length non-cell autonomously.

**Figure 4.**
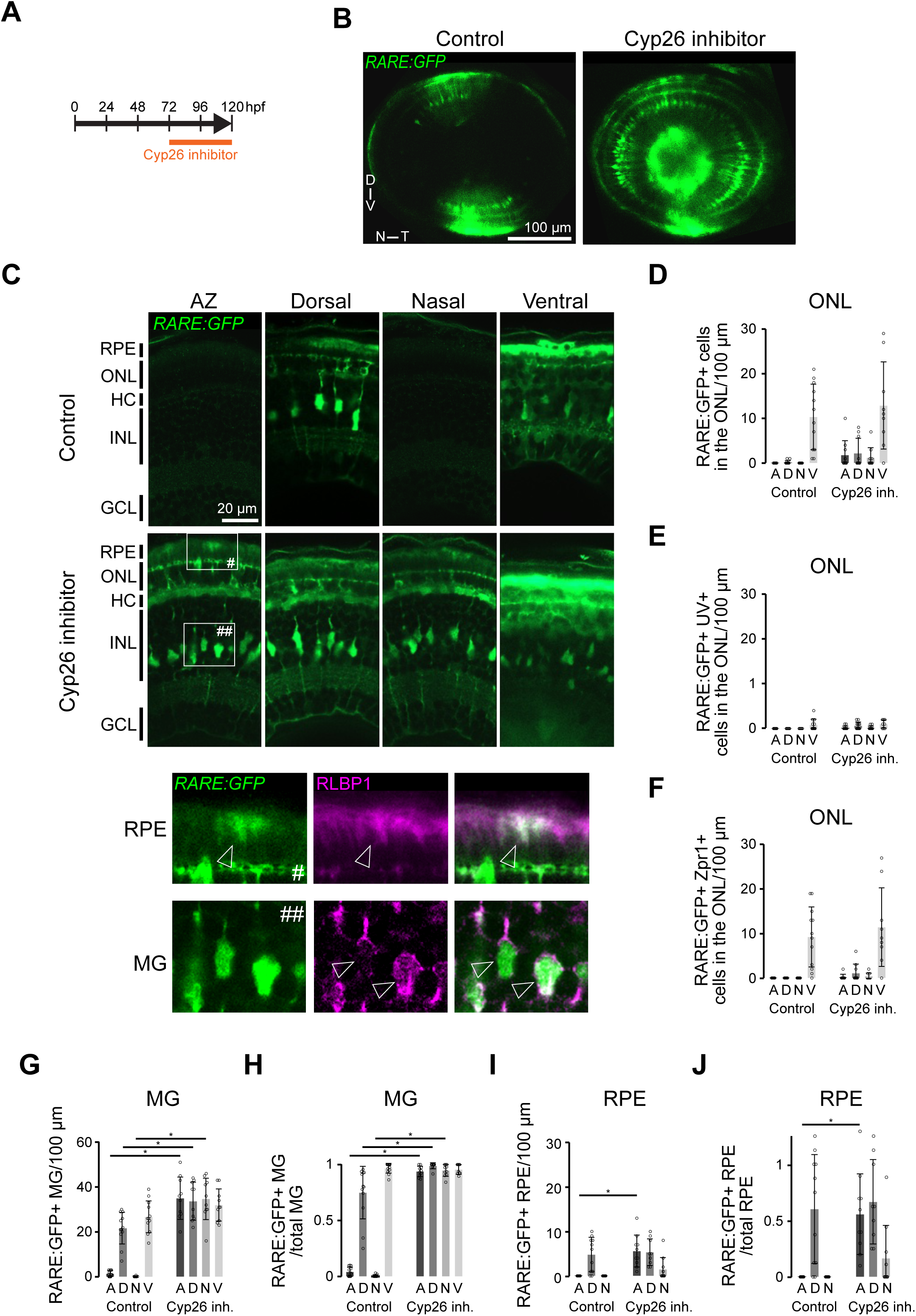
Cyp26 inhibition induces RA signalling predominantly in Müller glia and presumptive horizontal cells. **A**, Experimental timeline of drug application. **B**, Confocal images of the eye of *Tg(RARE:GFP)* zebrafish at 120 hpf. **C**, Example high-magnification images of different regions of those eyes. D, dorsal; N, nasal; T, temporal; V, ventral. GCL, ganglion cell layer; HC, horizontal cell; INL, inner nuclear layer; ONL, outer nuclear later; RPE, retinal pigment epithelium. Bottom images are high-magnification images of the rectangle regions in the ONL (#) and INL (##), respectively, co-labeled with an antibody against Rlbp1 to identify the retinal pigment epithelium (RPE, arrowheads) and Müller glia (MG, arrowheads). **D-F**, Densities of GFP-positive (**D,** n=12 (control) and n=10 (Cyp26 inh.)), GFP and UV opsin double-positive (**E**, n=12 (control) and n=12 (Cyp26 inh.)), and GFP and Zpr-1 double positive (**F**, n=12 (control) and n=10 (Cyp26 inh.)) cells in different regions of the ONL measured as the number of cells per 100 μm in a section. **G-J**, Densities and fraction of GFP-positive MG (**G**, n=12 (control) and n=10 (Cyp26 inh.); **H**, n=12 (control) and n=10 (Cyp26 inh.)) and RPE (**I**, n=12 (control) and n=10 (Cyp26 inh.); **J**, n=12 (control) and n=10 (Cyp26 inh.)) in the indicated regions of the eye. In all experiments in this figure, zebrafish were exposed to either vehicle or Cyp26 inhibitor from 72 to 120 hpf, and either imaged or fixed at 120 hpf. Bars: Mean, error bars: standard deviation. One-way ANOVA plus Tukey’s post-hoc test, *: p < 0.05 comparing the same region with or without Cyp26 inhibitor.

We then assessed in which cells RA signalling is activated following Cyp26 inhibition. Based on the cellular location and cell morphologies, we hypothesized that retinal pigment epithelial (RPE) cells, horizontal cells and Müller glia expressed GFP in R115866-treated retinas (Fig. 4B,C). To confirm that GFP-positive cells are Müller glia and RPE cells R115866- and vehicle-treated *RARE:GFP* transgenic eyes were immunolabelled with an antibody to Rlbp1, that identifies RPE in the outer retina and Müller glia in the INL (Fig. 4C). In vehicle controls, almost all the ventrally located Müller glia and a subset of Müller glia in the dorsal region were GFP-positive (Fig. 4G,H). In contrast, co-labelling of GFP and Rlbp1 was only observed occasionally in the AZ at the boundary to the ventral region and was absent in the nasal retina (Fig. 4G,H). Exposure to R115866 increased the number of GFP expressing Müller glia in the AZ, dorsal and nasal regions, resulting in almost all (>90%) Rlbp1-positive Müller glia express GFP in all regions (Fig. 4G,H). Some RPE cells in the ventral and dorsal retina but not in the AZ and the nasal regions expressed GFP in controls (Fig. 4I). Exposure to R115866 significantly increased GFP expression in a few RPE cells in the AZ (Fig. 4I). However, the proportion of GFP-positive RPE cells remained low (Fig. 4J).

We also confirmed that not only exogenous *RARE:GFP* reporter activity but also endogenous RA responsive genes are induced or suppressed following Cyp26 inhibition. HCR *in situ* hybridisation against *cyp26b1* and *cyp26c1*, which are two known RA-responsive genes [40] revealed that *cyp26b1* was upregulated (Supplementary Fig. 1), while *cyp26c1* expression was downregulated in both the ventro-nasal and dorso-temporal regions in R115866-exposed eyes (Supplementary Fig. 1). R115866-treatment in *RARE:GFP* transgenic zebrafish revealed that *cyp26b1* was expressed in GFP positive cells in the INL (Supplementary Fig. 1), which were Müller glia (Fig. 4C). This data suggests that: 1) to generate spatial RA signalling activity pattern, dorsal and ventral localised expression of RA synthesizing enzymes (*raldh1a2* and *raldh1a3*) are not sufficient but rather *cyp26* is critical for establishing the area of low RA signalling, and 2) Cyp26 inhibition overstimulates RA signalling in Müller glia and, to a lesser extent, in RPE cells, and almost none in UV cones.

### Region-specific UV cone OS shortening following RARα but not RARγ stimulation

To further confirm that the shortening of UV cone OS by Cyp26 inhibition is mediated by RA signalling we activated RA receptor using RA receptor alpha (RARα) and gamma (RARγ) agonists AM580 and BMS961, respectively, from 72 to 120 hpf. Exposure to the RARα agonist AM580 decreased the OS lengths across the regions measured (Fig. 5A,B). Relative OS length against dorsal OS revealed that the difference between the AZ and the dorsal, but not between the nasal and the dorsal, OSs was abolished, indicating that shortening of OS length is not uniform across the regions but rather enhanced in the AZ (Fig. 5A,C). In contrast, exposure to the RARγ agonist BMS691 did not significantly affect the length of UV cone OSs or the d/AZ and d/n ratios (Supplementary Fig. 2). In agreement, GFP expression was upregulated broadly throughout the entire eye when *RARE:GFP* transgenic zebrafish were exposed to the RARα agonist, but not by RARγ agonist (Fig. 6A). This data suggests that RARα are the predominant RA receptors acting in the retina. Moreover, activation of RA signalling similarly shortens UV cone OS as seen after Cyp26 inhibition, indicating that the effect of Cyp26 inhibition on the UV cone OS length is mediated by RA signalling.

**Figure 5.**
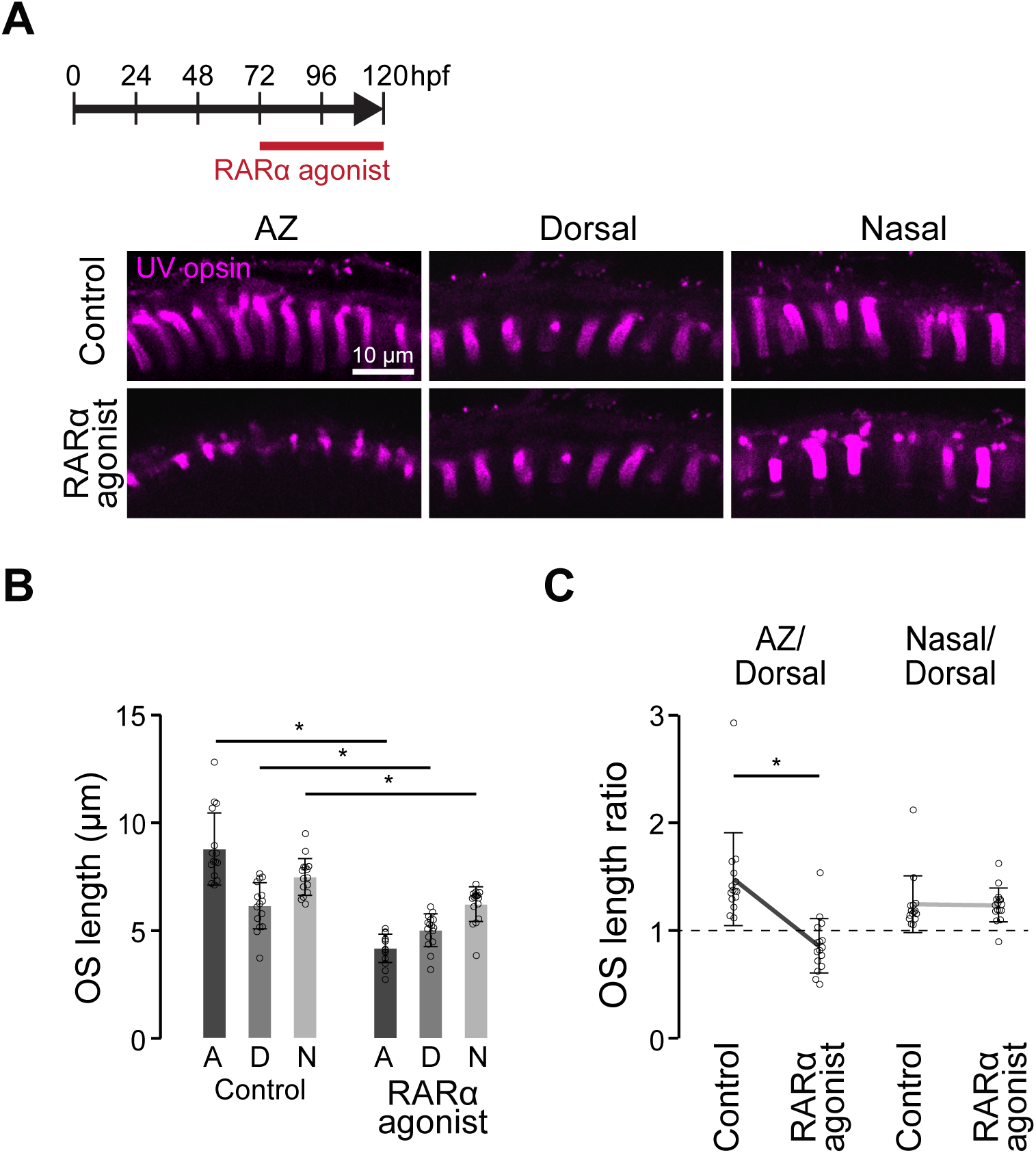
RARα mediated Retinoic Acid signalling shortens OS in the AZ. **A**, Experimental timeline of drug application and confocal images of the UV opsin immnostaining in the photoreceptor layer in the AZ, dorsal and nasal retinas labelled at 120 hpf. Zebrafish were treated with control vehicle or a RARα agonist, AM580, from 72 to 120 hpf. **B**, UV cone OS length in the AZ, dorsal and nasal regions of the retina at 120 hpf. **C**, Comparisons of the ratios of the UV cone OS length of the AZ or nasal regions against the dorsal region with and without AM580. Bars and lines: mean. Error bars: standard deviation. Open circles: data from individual fish. n=15 (Control) and 17 (RARα agonist). One-way ANOVA plus Tukey’s post-hoc test, *: *p* < 0.05.

**Figure 6.**
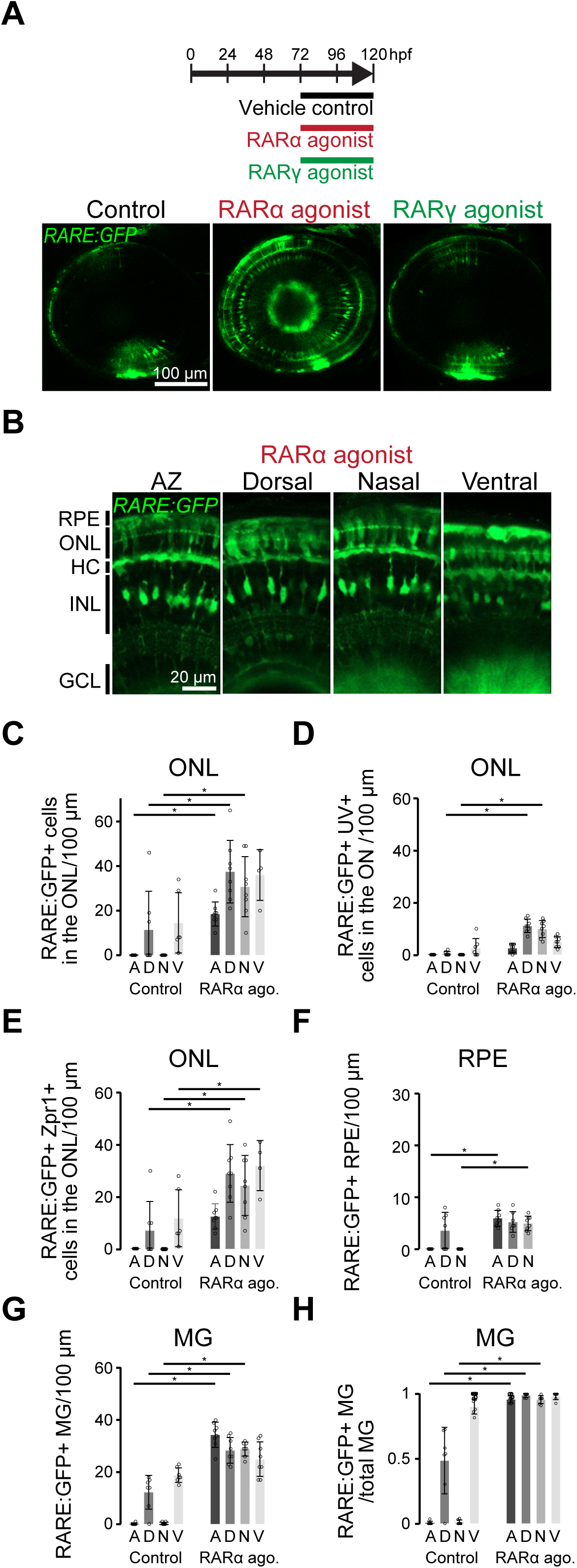
RARα activation induces RA signalling in specific cell-types similar to Cyp26 inhibition. **A**, Experimental timeline of drug applications and confocal images of the eye of *Tg(RARE:GFP)* zebrafish at 120 hpf after treatment with vehicle control, RARα and RARγ agonists. D, dorsal; N, nasal; T, temporal; V, ventral. **B**, High-magnification images of different regions of the RARα agonist-treated *Tg(RARE:GFP)* eyes. GCL, ganglion cell layer; HC, horizontal cell; INL, inner nuclear layer; ONL, outer nuclear later; RPE, retinal pigment epithelium. **C-E**, Densities of GFP-positive (**C,** n=8 (control) and n=7 (RARα agonist)), GFP and UV opsin double-positive (**D**, n=7 (control) and n=8 (RARα agonist)), and GFP and Zpr-1 double positive (**E**, n=7 (control) and n=8 (RARα agonist)) cells in different regions of the ONL measured as the number of cells per 100 μm in a section. **F**, Densities of GFP-positive RPE (n=8 (control) and n=8 (RARα agonist). **G**,**H**, Densities (**G**, n=7 (control) and n=8 (RARα agonist)) and fraction (**H**, n=7 (control) and n=7 (RARα agonist)) of GFP-positive MG in the indicated regions of the eye. Bars: Mean, error bars: standard deviation. One-way ANOVA plus Tukey’s post-hoc test, *: p < 0.05 comparing the same region with or without RARα agonist.

We further examined whether RARα agonist activates RA signalling in the same cell-type as observed in Cyp26 inhibition. We co-labelled *RARE:GFP* transgenic retinas with cell type specific markers after exposure to RARα agonist. In the ONL, RARα agonist significantly increased the number of GFP-positive cells in the AZ, dorsal, and nasal retinal regions relative to controls (Fig. 6B,C). The number of GFP-positive UV cones remained low (Fig. 6D), while red and green cones account for the majority of GFP-positive cells in the ONL (Fig. 6E). In particular, GFP-positive UV cones were unchanged in the AZ, while they significantly increased in the dorsal and nasal regions in RARα agonist-treated zebrafish. (Fig. 6E).

We next assessed whether RA signalling is activated in Müller glia and RPE cells by RARα agonist as seen by Cyp26 inhibition. We observed an increase in GFP-positive RPE cells in the AZ and the nasal retina where normally RPE cells are GFP-negative (Fig. 6F). Similarly, we observed that the number of GFP-positive Müller glia increased in the AZ, dorsal, and nasal regions where typically only some or very few Müller glia express GFP (Fig. 6G). RA signalling was activated in almost all Müller glia across the regions following exposure to RARα agonist (Fig. 6H). Thus, cell type specificity of RA signalling activation and region-specific OS shortening by RARα agonist are consistent with the observations in Cyp26 inhibition condition.

### OS shortening leads to reduced light sensitivity in UV cones

To determine if the shortening of OS by RA signalling reduces light sensitivity, we recorded UV cone calcium responses to varying intensities of light stimulations. We used the transgenic line *Tg(opn1sw1:sypb-GCaMPf)^uss101^*, which we previously used to characterize regional differences in UV cone light sensitivity in the eye [3]. In this line, synaptically tagged fluorescent calcium biosensor SyGCaMP6f is expressed in UV cones (Fig. 7A). Larvae at 5 dpf, treated with either vehicle control or RARα agonist from 3 dpf until imaging, were imaged under a two-photon microscope at 16.7 Hz (256 x 60 pixels). This setup allowed us to simultaneously capture 10-20 UV cone synaptic calcium responses during light stimulation. We presented light flashes of varying duration from darkness (Fig. 7B) and measured the amplitudes of light-evoked synaptic calcium signals. As expected, UV cones responded to light with a reduction of synaptic calcium. The amplitude of this reduction increased with longer light flash durations. We previously reported that UV cones in the AZ exhibit larger calcium responses than dorsal UV cones, corresponding to the longer OS in the AZ UV cones [3]. We confirmed the same result in control larvae; the light sensitivity is higher in AZ UV cones than dorsal UV cones (Fig. 7B,C). However, we found that overall light sensitivity was reduced by RARα agonist treatment in both AZ and dorsal UV cones, and that the regional difference was abolished (Fig. 7B,C). These results indicate that changes in the OS length by RA signalling accompanies with changes in light sensitivity.

**Figure 7.**
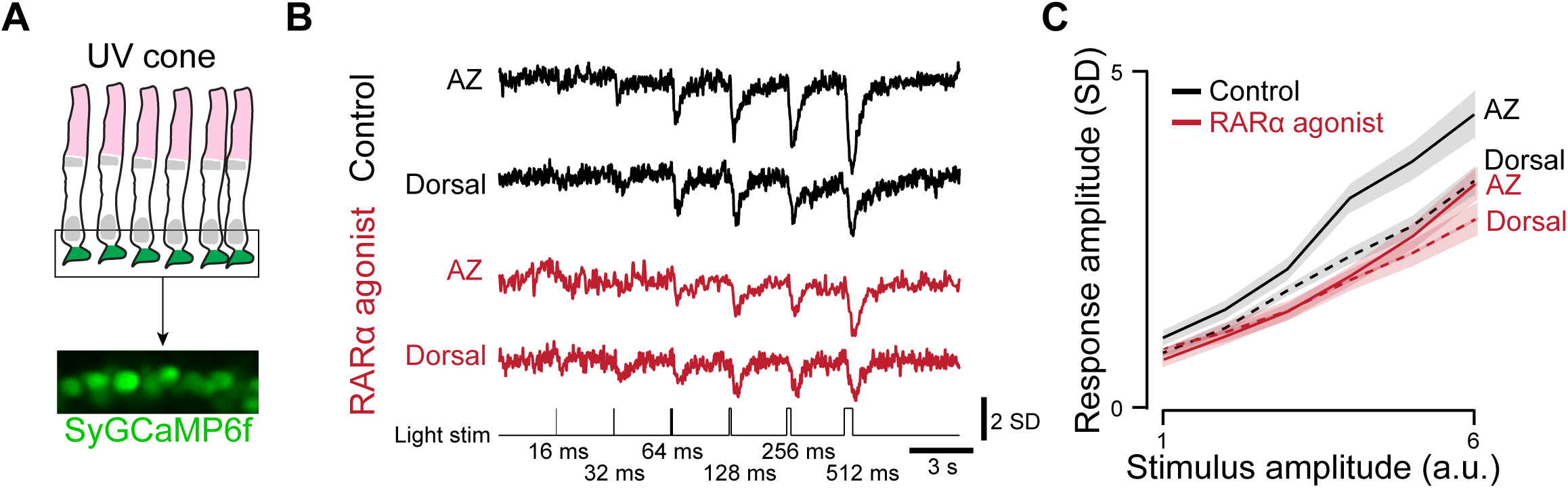
Light sensitivity is reduced in UV cones with shorter OSs after RARα agonist treatment. **A**, Schematic and a 2-photon image of synaptically-targeted calcium sensor, SyGCaMP6f, in UV cones. **B**, Example single cone 2-photon calcium responses to varying duration of UV light flashes (1.5 × 10^4^ photon/s/μm^2^). Light stim: UV light stimulation. **C**, Mean peak response amplitudes. Shadings indicate 95% confidence intervals. n= 5 and 4 fish for control and RARα agonist, respectively. n = 141, 111, 109, 63 UV cones for AZ (control), Dorsal (control), AZ (RARα agonist), Dorsal (RARα agonist), respectively.

### Müller glia are morphologically specialised in the AZ and required for PR specialisation

The expression of *RARE*-driven GFP in Müller glia, horizontal cells and RPE cells suggests that one or multiple of these cells may exert a non-cell autonomous effect on UV cones, thereby modulating their OS growth. Müller glia possess apical microvilli that are in close contact with photoreceptor inner segments and OSs [41,42] (Fig. 8A). Furthermore, MG apical microvilli have been implicated in maintaining the morphological structure of photoreceptor OSs [43–45]. RPEs are also known to be critical for maintaining OS structures and functions [46,47]. However, we observed that a relatively small fraction (∼50%) of the RPE activates RA signalling in the AZ following Cyp26 antagonist treatment, whereas RA signalling is active in almost all Müller glia in the same condition (Fig. 4H,J). This positions Müller glia as a prime candidate to exert a non-cell autonomous effect on UV cones following stimulation of RA signalling.

**Figure 8.**
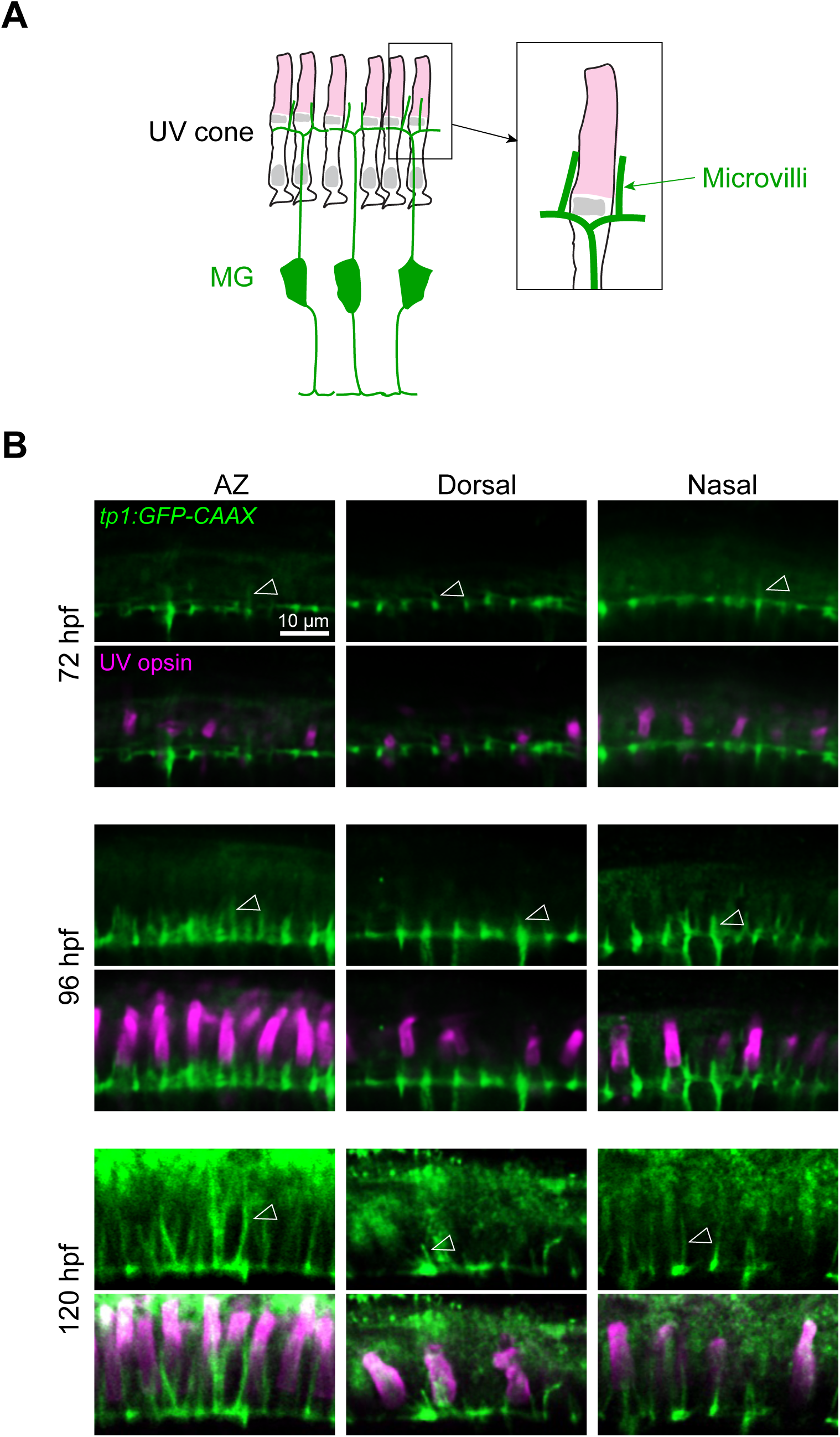
Müller glia apical microvilli develop in a region-specific manner in a similar timeframe as UV cone outer segments. **A**, Schematic of a cross-section of a larval zebrafish retina (left) displaying the apical microvilli of Müller glia (MG, green) in close contact with UV cone outer segments (light magenta). **B**, Confocal images of the photoreceptor OS layer in *Tg(tp1:GFP-CAAX)* zebrafish labelled with anti-UV cone opsin antibody in the AZ, dorsal and nasal retina at 72, 96 and 120 hpf. Arrowheads indicate apical microvilli of Müller glia.

It is currently unknown whether Müller glia apical microvilli display region-specific differences and whether they play a role in photoreceptor OS development. Thus, we first assessed the development of Müller glia apical microvilli using *Tg(TP1:GFP-CAAX)* zebrafish that express membrane-bound GFP in Müller glia, which allows the visualisation of small processes such as apical microvilli [29]. At 72 hpf, 12 hours after Müller glia are born [35], a few very short apical microvilli were observed in the nasal retina and AZ. In contrast, apical microvilli were absent in the dorsal retina (Fig. 8B). Subsequently at 96 hpf, a regular pattern of short microvilli was observed in the AZ, nasal and dorsal regions (Fig. 8B). The Müller glia apical microvilli further extended by 120 hpf, displaying long microvilli in the AZ and nasal regions in contrast to short microvilli in the dorsal retina (Fig. 8B). These observations demonstrate that Müller glia display region specific morphologies, with apical microvilli length corresponding to the UV cone OS length.

To investigate whether Müller glia are necessary to mediate the RA-induced shortening of UV cones we prevented Müller glia genesis in the developing retina by applying the γ-secretase inhibitor DAPT from 45 hpf and alongside either the vehicle control or AM580 from 72 to 120 hpf (Fig. 9A,B, Supplementary Fig. 4). UV cone OSs were significantly shorter in all analysed regions (AZ, nasal, dorsal) in DAPT-treated embryos that lack Müller glia (Fig. 9C,D). Furthermore, the ratios of OS length between the AZ or the nasal, and the dorsal regions were similarly reduced to approximately 1, indicating that regional specialisation was lost when Müller glia are absent (Fig. 9C-E). Applying RARα agonist in the absence of Müller glia, i.e. to DAPT-treated retinas, did not further reduce OS length compared to DAPT-treated retinas alone (Fig. 9C-E). To test whether DAPT still prevent the regional OS growth in the presence of Müller glia, we applied DAPT from 72 hpf when Müller glia have been produced but prior to UV cone OS growth (Fig. 9F,G). We found that the regional specialisation of UV cones was unaffected by DAPT-treatment based on the UV cone OS length and the ratio analysis of the OS length between different retinal regions (Fig. 9H,I), indicating that blocking γ-secretase activity during OS growth does not result in UV cone OS shortening. Taken together, this data demonstrates that Müller glia are required for OS specialisation and supports the hypothesis that Müller glia modulate UV cone OS growth via RA signalling in a non-cell autonomous manner.

**Figure 9.**
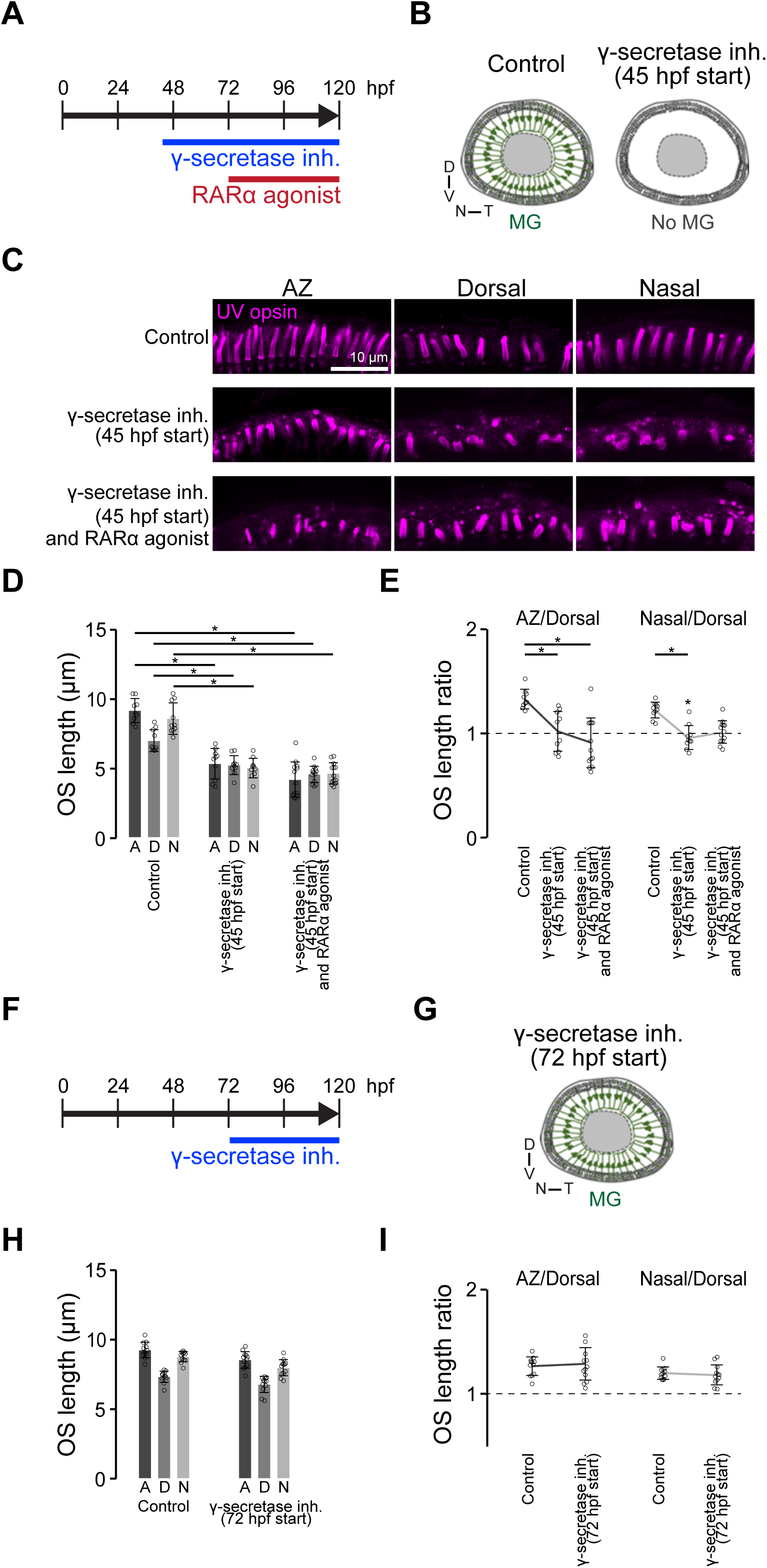
Absence of Müller glia shortens UV cone OS in the entire retina. **A**,**B**, Timeline of γ-secretase inhibitor (inh.) (from 45 hpf) and RARα agonist (from 72 hpf) exposures (**A**), and schematics of eyes displaying the presence and absence of Müller glia (MG, green) following eFont fixedxposure of zebrafish to control vehicle or the γ-secretase inhibitor, respectively, from 45 to 120 hpf (**B**). **C**, Confocal images of the UV cone OS stained with anti-UV opsin antibody in the AZ, dorsal and nasal retina at 120 hpf following pharmacological treatments as indicated in (**A**). **D**,**E**, UV cone OS length (**D**) and the ratios of the UV cone OS length (the AZ or nasal regions against dorsal region) (**E**) following pharmacological treatments as indicated in (**A**). n=10 (control); n=10 (γ-secretase inhibitor); n=13 (γ-secretase inhibitor and RARγ agonists). **F**,**G**, Timeline of γ-secretase inhibitor treatment (from 72 hpf after Müller glia genesis) and schematic depicting the resulting eye with Müller glia present (green). **H**,**I**, UV cone OS length (**H**) and the ratios of the UV cone OS length (the AZ or nasal regions against dorsal region) (**I**) following pharmacological treatments as indicated in (**F**). n=12 (control); n=12 (γ-secretase inhibitor). Bars and Lines: mean. Error bars: standard deviation. Open circles: data from individual fish. One-way ANOVA plus Tukey’s post-hoc test, *: *p* < 0.05.

## Discussion

By focusing on UV cones in zebrafish, we identify glia as a regional modifier of RA signalling to regulate regional variations in cone OS length and function. We further found that the RA’s effects on cone OS length is non-cell autonomous, likely mediated by Müller glia. Taken together, these findings highlight the importance of glia in establishing regional specializations of neurons.

The roles of RA signalling in early eye development have been well characterized, including in photoreceptor differentiation [48–53]. Across vertebrate species, RA promotes rod differentiation and modulates cone subtype specification. In zebrafish, RA signalling accelerates rod and red cone differentiation while supressing blue and UV cone opsin expression [48,49]. In human retinal organoids, early RA exposure promotes rod and M cone fates [50–53]. Furthermore, studies in chicken demonstrated that this cell type specification by RA signalling create regional differences in cell type composition in the retina [20]. In chicken, photoreceptors in the high-acuity area (HAA; an analogous structure of the human fovea or the zebrafish AZ) is composed exclusively of cones. There, localised *cyp26* expression generates a region of low RA signalling, which causes lack of rod genesis. These findings indicate that local suppression of RA signalling contributes to generating HAA specific photoreceptor composition during early cell differentiation. However, despite extensive evidence for RA signalling regulating cell fate during development, its role in cellular specialization after cell differentiation has been less explored. Our findings in this study demonstrate a novel role of RA signalling beyond fate specification, regulating regional functional specialisations on photoreceptors. In humans, Müller glia appears as early as 7 weeks in the fovea, before the formation of neuronal specialisations and circuit development, including cone outer segments, required for high acuity vision [54]. Furthermore, Cyp26a1 is specifically expressed in the macular Müller glia [27]. Taken together, we speculate that the mechanisms of Muller glia dependent HAA specializations we found in zebrafish may exist in the human macular.

It has been proposed that longer cone OSs in the primate fovea arise passively from tight cone packing, leading to physical elongation. In macaques, cone OSs are longer in the fovea than the periphery, but the total volume of cone OS does not differ between these two regions [55], supporting that the elongation of cone OS in the fovea is not due to enhanced OS growth. While such passive mechanical factor may still contribute, our findings reveal that OS length is also actively regulated by molecular signalling. Supporting this, foveal cone OS layer thickness can change depending on conditions.

Cone OS layer becomes thinner in vitamin A deficiency patients. However, in some patients, vitamin A supplementation restores cone OS layer thickness [56], suggesting that cone OS length is actively regulated.

Depending on species, HAAs, including the fovea and the zebrafish AZ, exhibits multiple forms of specialisations beyond cone OS length. As mentioned above, in the chicken retina, rod exclusion requires locally reduced RA signalling [20]. This mechanism also exists in zebrafish. Rods are enriched in dorsal and ventral retina, which correspond to the areas with shorter cone OSs. RA treatment during early development expands the rod enriched areas [48]. However, whether RA signalling also regulates differentiation of other HAA-specific cell types or later aspects of their functional specialisation remains to be determined. Late developmental RA exposure in zebrafish increases rhodopsin and red opsin gene expression while supressing blue and UV opsin expressions [49]. This effect is not due to cell fate conversion [49], hinting that RA also regulate functional maturation of regional specialisations in other cones.

Manipulations of Cyp26 and RA signalling were conducted by systemic treatment of zebrafish with pharmacological drugs, rather than using *cyp26a1* and *c1* mutants, to investigate the specific role of Cyp26 after cone genesis. In fact, we found that the retinal development is severely disrupted in cyp26a1 and c1 double mutant (unpublished). In contrast, we found that the gross morphology of the retina is preserved after pharmacological manipulation of RA signaling from 3-5 dpf (for example, Fig. 4A). The limitation of systemic pharmacological treatments is that we cannot exclude the possibility that the changes in UV cone OS length were due to activation of RA signalling outside of the eye, potentially through release of secretary factors outside of the eye after RA signalling activation and the factors reach the eye and modulate cone OS length. However, this scenario does not explain why cone OS length are regionally different in normal condition and why Cyp26 inhibition specifically shortens cone OS in the AZ than other regions. Our observation that the RA signalling activity pattern within the eye match the variation in cone OS length strongly suggests that RA signalling within the eye locally regulate cone OS length.

How RA signalling in Müller glia regulate cone OS length remains an open question. Upon RA activation either by Cyp26 inhibition or by RARα agonist, the majority of Müller glia throughout the entire retina upregulate *RARE:GFP* expression as well as the RA-responsive gene, *cyp26b1*. However, the downstream effects on cone OS growth are most pronounced in the AZ (Fig. 3E). This suggests that RA-induced effectors that restrict OS growth in the AZ must be specifically up or down-regulated within this region and will need to be identified in the future.

Glia are increasingly being appreciated for roles beyond metabolic and physical support for neurons. They act as active regulators of morphogen gradients in the developing CNS to promote neuronal proliferation, differentiation and axon guidance [57–60]. Here we show that regional regulation of morphogen activity by glia controls specialisation for high-acuity visual function in photoreceptor neurons. This raises the tantalising possibility that other glia may also regulate neuronal specialisation and function by integrating morphogen signals during development. There is emerging evidence suggesting specific roles in astrocyte-neuron interactions, including astrocytes being identified as a source for RA in the brain [61], secreted RA from astrocytes maintains neuronal structure in vitro [62], and has been implicated under pathological conditions in gliomas and Alzheimer’s disease [62,63]. is a common feature in many neurological and psychiatric disorders. A recent study has found that S100A6 acts as a glial derived morphogen that impacts neurite development [64]. However, the role of glia in the CNS in the regulation of morphogen activity in development to fine tune or impact neuronal function remains unclear.

## ACKNOWLEDGEMENTS

We thank Yi Jiang for critical reading of the manuscript. We thank Aanandita Kothurkar for technical and cloning assistance. This study was supported by a Macular Society grant (14225), Alcon Research Institute (AW00013337), Research to Prevent Blindness and NEI (R01EY036090) to TY, a BBSRC David Phillips Fellowship (BB/S010386/1) to R.B.M., and Moorfields Eye Charity Springboard award (GR001388) to T.Y. and R.B.M.

## AUTHOR CONTRIBUTIONS

T.Y. and R.B.M. conceived the study and M.L., T.Y., and R.B.M. designed the experiments. R.L. conducted imaging of OS in treatment groups and performed OS quantifications. M.S. performed the measurements of the correlation between RA signalling and cone OS length with inputs from T.Y. and T.Y. performed calcium imaging experiments. M.L. performed all the rest of data collection with inputs from T.Y. and R.B.M. M.L., T.Y., and R.B.M. wrote the manuscript.

---

**Supplementary Figure 1.**
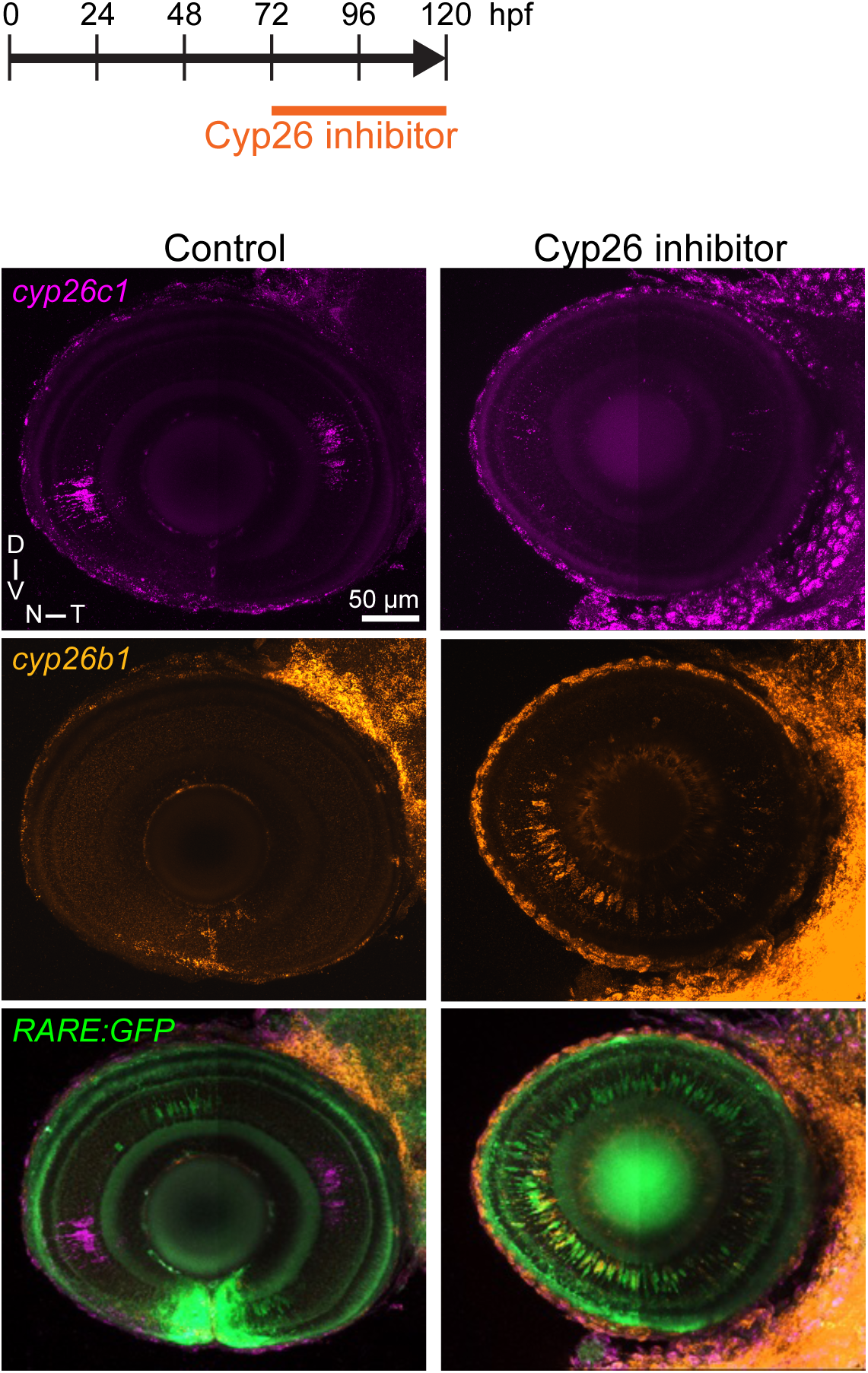
Cyp26 inhibition induces differential expression of retinoic acid target/responsive genes. Experimental timeline of drug application and confocal images of HCR *in situ* hybridisation against *cyp26c1* (magenta) and *cyp26b1* (orange) in *Tg(RARE:GFP)* (green) transgenic zebrafish at 120 hpf. Zebrafish were treated with or without Cyp26 inhibitor from 72 to 120 hpf.

**Supplementary Figure 2.**
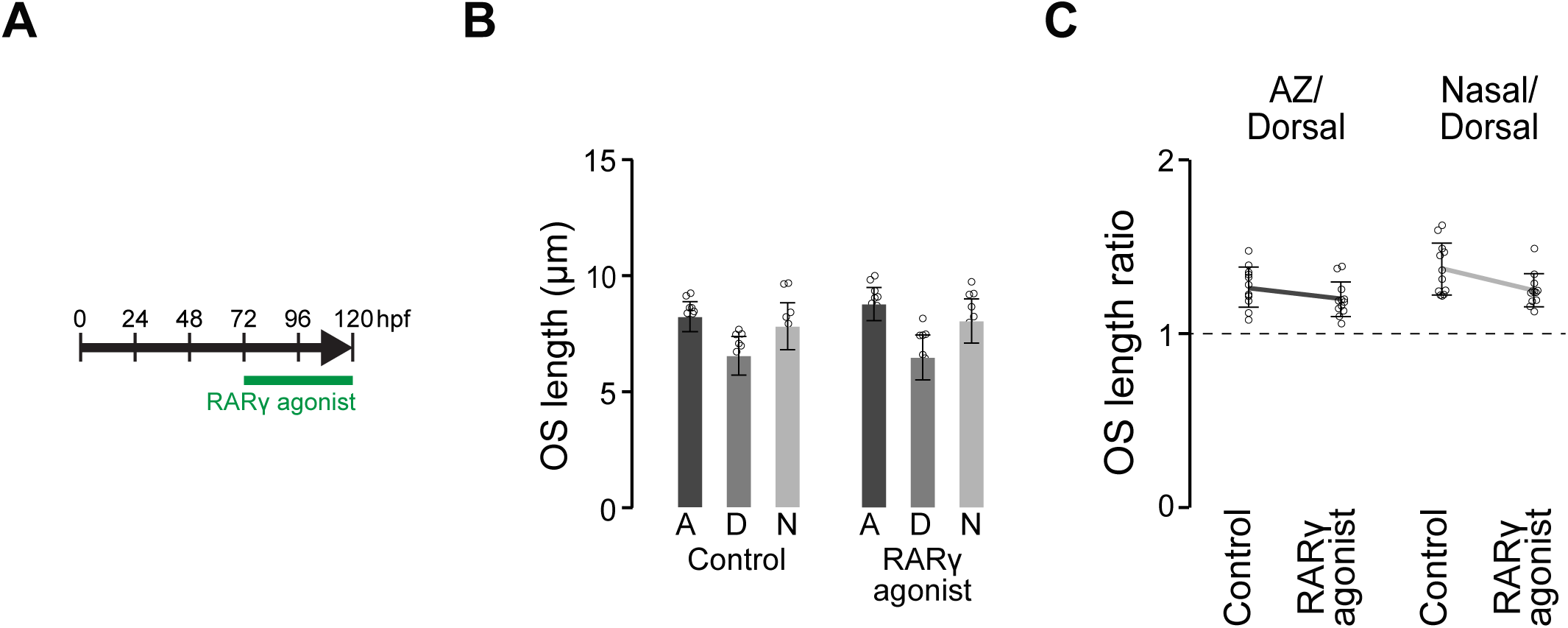
RARγ activation does not change UV cone OS length. A, Experimental timeline of drug application. **B**,**C**, UV cone OS length (**B**) and the ratios of the UV cone OS length (the AZ or nasal regions against dorsal region) (**C**) at 120 hpf following RARγ agonist treatment from 72 to 120 hpf. n=12 (control); n=12 (RARγ agonist). Bars and Lines: mean. Error bars: standard deviation. Open circles: data from individual fish.

**Supplementary Figure 3:**
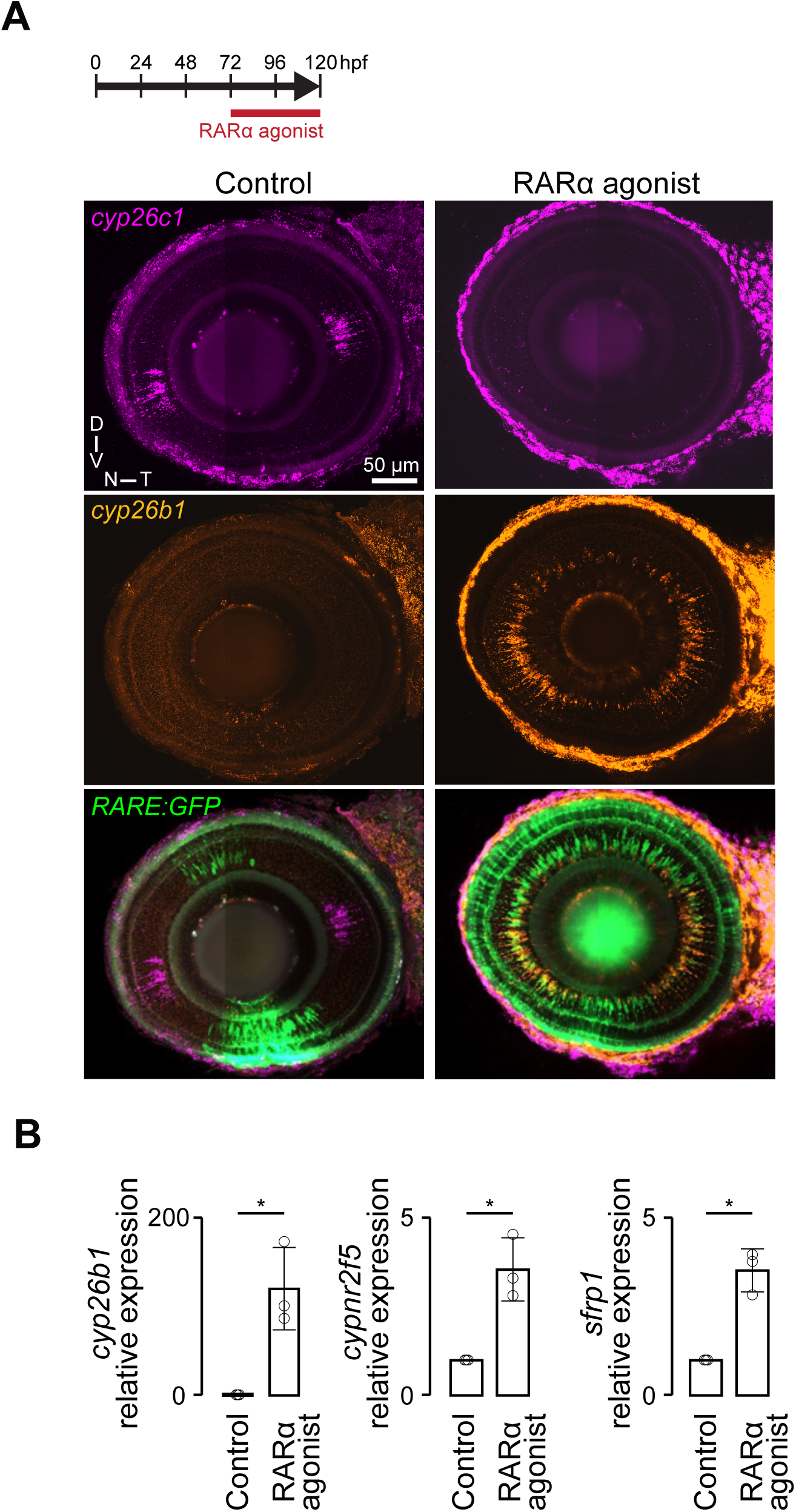
RARα activation induces differential expression of retinoic acid target/responsive genes, similar to Cyp26 inhibition. **A**, Experimental timeline of drug application and confocal images of HCR *in situ* hybridisation against *cyp26c1* (magenta) and *cyp26b1* (orange) in *Tg(RARE:GFP)* (green) transgenic zebrafish at 120 hpf. Zebrafish were treated with or without RARα agonist from 72 to 120 hpf. **B**, Relative mRNA expression changes for retinoic acid target/response genes *cyp26b1*, *nr2f5* and *sfrp1a* between controls and RARα agonist-exposed samples. Mean ± S.E., n = 3, Student’s t-test, *p < 0.05.

**Supplementary Figure 4:**
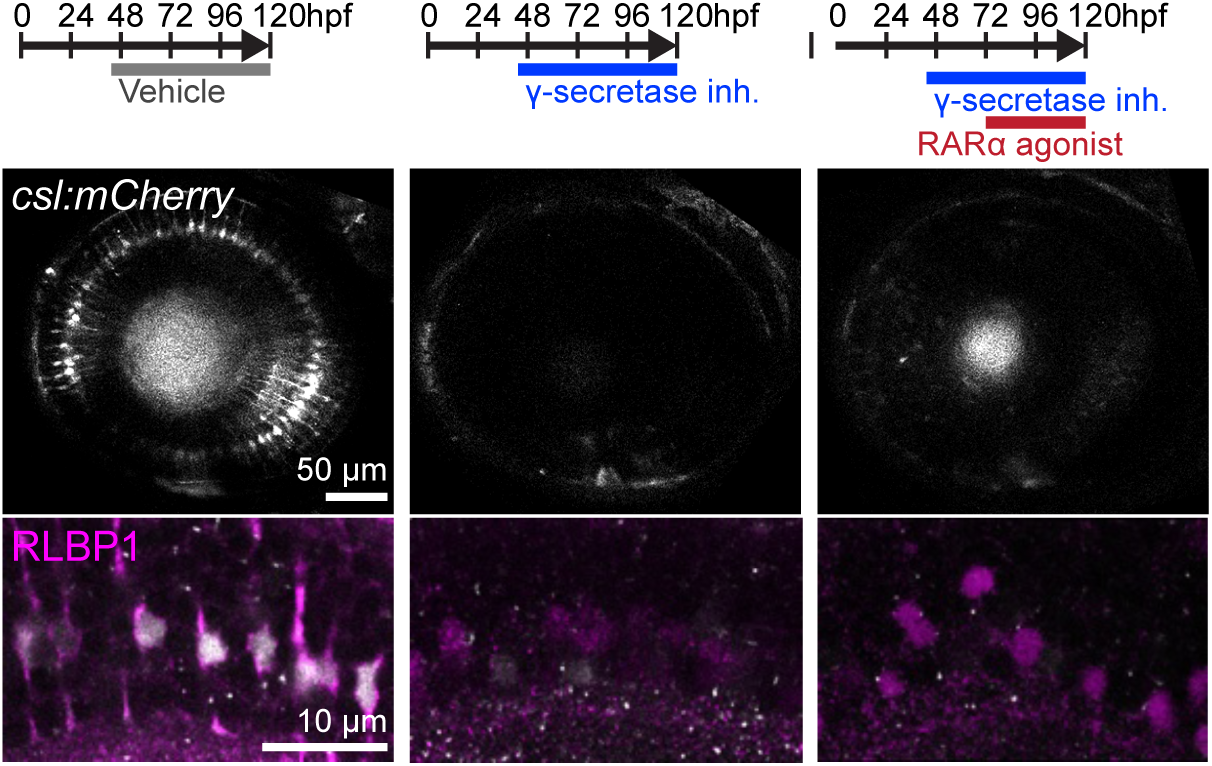
Müller glia are absent in γ-secretase inhibitor treated retinas. Experimental timeline of drug application and confocal images of the *Tg(csl:mCherry)* transgenic eyes at 96 hpf (top row) and high-magnification images in the inner nuclear layer in the AZ at 120 hpf immunolabelled with the Müller glial marker Rlbp1 (magenta, bottom row). Fish were exposed to γ-secretase inhibitor, DAPT or control vehicle from 45 hpf. At 72 hpf, the fish were additionally exposed either to the RARα agonist, AM580 or control vehicle. Note that some amacrine cells become RLBP1 positive in the γ-secretase inhibitor plus RARα agonist condition.

**Supplementary Figure 5:**
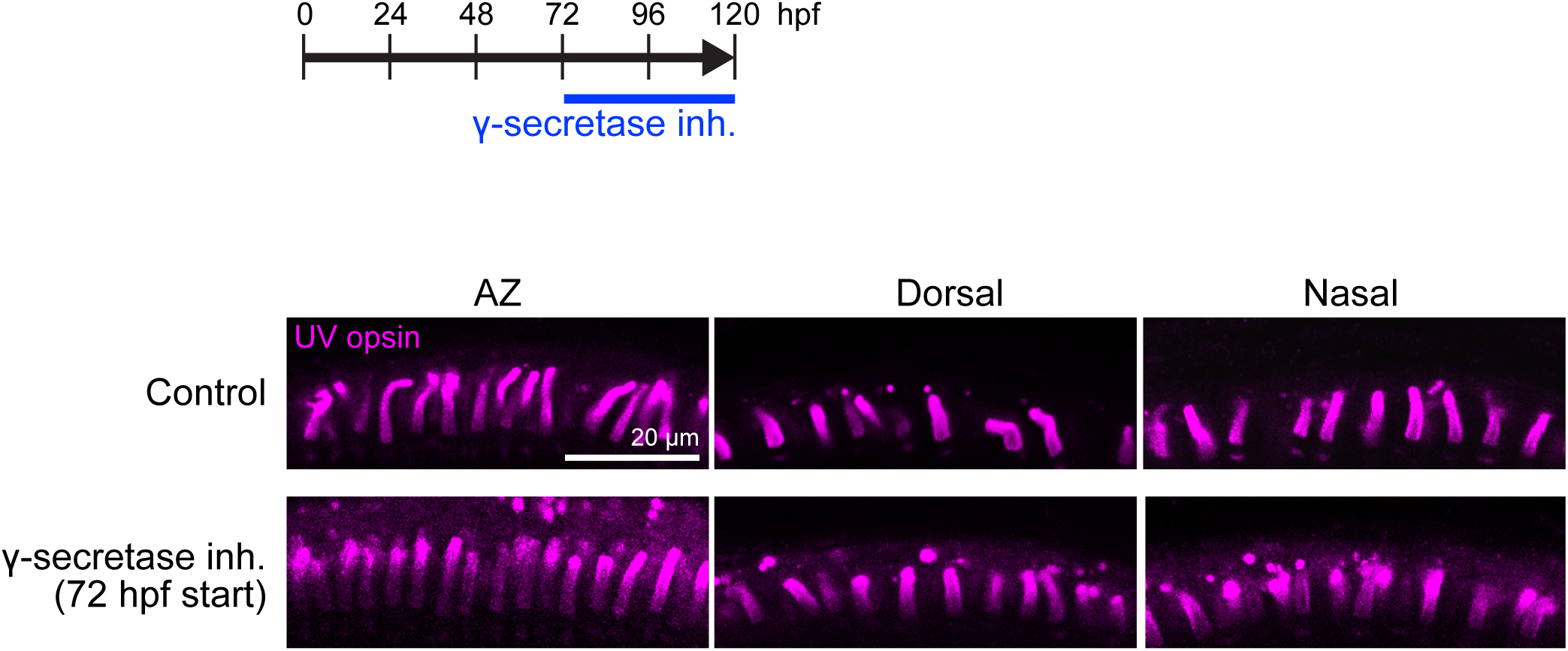
Inhibition of γ-secretase from 72 hpf does not change UV cone OS length. Timeline of γ-secretase inhibitor treatment (from 72 to 120 hpf) and confocal images of the photoreceptor layer immunolabelled with anti-UV opsin antibody in the AZ, dorsal and nasal retina at 120 hpf.

